# Highly Pathogenic Avian Influenza A(H5N1) Caused Mass Death among Black-legged Kittiwakes (*Rissa tridactyla*) in Norway, 2023

**DOI:** 10.1101/2025.05.23.655725

**Authors:** Grim Rømo, Caroline Piercey Åkesson, Tone Kristin Reiertsen, Johanna Hol Fosse, Cathrine Arnason Bøe, Lars Austbø, Johan Åkerstedt, Maryam Saghafian, Morten Helberg, Olav Hungnes, Britt Gjerset, Silje Granstad, Gørill Hogseth, Siri Løtvedt, Anne Døsen, Ragnhild Tønnessen

**Author notes:** Corresponding author: Grim Rømo.

## Abstract

In 2023, highly pathogenic avian influenza (HPAI) heavily affected gulls in Europe. In July, a mass mortality event was reported in the Black-legged Kittiwake (*Rissa tridactyla*) breeding colony at Ekkerøy in Northern Norway. The cause was confirmed to be infection with the HPAI H5N1 clade 2.3.4.4b virus, genotype EA-2022-BB. We describe the outbreak in Kittiwakes, including pathological and virological investigations, and discuss the management and zoonotic potential. With more than 15,000 dead birds reported, we estimate that the outbreak caused a reduction in the Kittiwake population at Ekkerøy of at least 50%. Diseased birds exhibited neurological signs. Necropsy of ten birds revealed a peracute fatal systemic disease, with severe lesions in the brain and pancreas co-localizing with the presence of viral RNA and antigen. Vascular expression of α2,3-linked sialic acids and viral RNA/antigen may reflect hematogenous virus spread. Further studies should investigate the long-term impact of HPAI on Kittiwake populations.

## Background

From autumn 2020 to 2023, three major highly pathogenic avian influenza (HPAI) epizootics caused by H5Nx clade 2.3.4.4b viruses hit Europe, causing suffering and death to millions of wild and domestic birds (1). During this period, H5N1 became the predominant avian influenza virus (AIV) subtype and spread with migratory birds across North and South America, Africa, and the Antarctic region, causing a panzootic (2, 3). Sporadic spillover has caused HPAI infection in several wild and domestic mammals (1), with outbreaks among farmed fur animals in Europe (4, 5), and massive mortality among marine mammals in South America (6). HPAI H5N1 virus also caused a multistate outbreak in dairy cattle in the USA and spillover from captive birds to a sheep in England (7, 8). So far, HPAI H5N1 viruses have only sporadically caused disease in humans (9). However, the expanding host range of HPAI H5N1 clade 2.3.4.4b viruses with increased circulation among domestic mammals elevates the zoonotic threat.

Several genotypes of H5N1 have circulated in birds in Europe (10). From February to October 2023, the Eurasian BB genotype (H5N1-A/Herring_gull/France/22P015977/2022-like), also known as EA-2022-BB, became predominant in Europe (11). This genotype contains three H13 virus-derived gene segments. Since gulls are the primary hosts of low pathogenic (LPAI) H13 viruses, this likely contributes to the virus’s adaptation to these species (12).

In spring 2023, mortality among terns and gulls increased dramatically in coastal breeding colonies, especially in the Wadden Sea region (Netherlands, Germany and Denmark), Great Britain and in Latvia (11). In Norway, the EA-2022-BB genotype was first detected in a dead Herring gull (*Larus argentatus*) in April 2023, but during summer Black-legged Kittiwake (*Rissa tridactyla*) became the most severely affected species.

Kittiwakes migrate along the eastern parts of the Atlantic. Most birds reside in the western part of the Atlantic Ocean during winter, but some also stay in the eastern parts before returning to their colonies. From the age of three to five years, they breed in colonies on steep cliffs along coastlines in the boreal and arctic zones of the northern hemisphere. The Kittiwake has status as vulnerable in the IUCN Red List of Threatened Species and is considered endangered on the Norwegian Red List for Species (13). In 2015, the total Kittiwake population on the Norwegian mainland was estimated to be 87,000 breeding pairs, but has since been declining (14). The colony on Ekkerøy (70°4’ N, 30°7’ E) is the largest on mainland Norway. The most recent systematic estimate of the Ekkerøy colony was conducted in 2012, counting 17,000 breeding pairs (15). Ekkerøy is located in Vadsø municipality, Northern Norway, and is known as a popular site for birdwatching. Vadsø is the county capital of Finnmark. It has 5,800 inhabitants and a 78.4 km coastline.

Previous studies have detected AIVs in Kittiwakes in Norway (16) and Canada (17). Antibodies against AIVs, including H13 and H16, have been detected in Kittiwakes in mainland Norway (16), Svalbard (18), and Scotland (19), suggesting that these viruses circulate in Kittiwakes of the North Atlantic. There is limited knowledge about the occurrence of HPAI H5 viruses in this species (20). No studies on the pathology or receptor distribution of AIVs in Kittiwakes have been published, and the impact of AIV infection on Kittiwake colonies is still largely unknown.

In July 2023, a mass mortality event hit the breeding colony of Kittiwakes at Ekkerøy. We describe clinical signs, pathology, and virus receptor distribution in tissues. Furthermore, we characterize the causative H5N1 virus and its zoonotic potential, discuss the mitigation efforts initiated by the authorities, and provide recommendations for management of future outbreaks in colonies.

## Material and Methods

This section gives a brief overview of material and methods used, with further details provided in Appendix 1.

Confirmed cases of HPAI virus detections in Kittiwakes in Norway from May 1 to July 31, 2023, were extracted from surveillance data from the Norwegian Veterinary Institute (NVI). Kittiwake mortality data were obtained from the County Governor of Troms and Finnmark. To estimate the number of Kittiwake pairs breeding at Ekkerøy prior to the outbreak, an ad hoc survey of occupied nests was conducted by the Norwegian Institute of Nature Research and the Norwegian Nature Inspectorate on July 27, 2023.

Clinical observations of Kittiwakes were mainly conducted at the breeding colony at Ekkerøy and at the river outlet of Storelva, 3.2 km north of the colony, where a high number of birds gathered. To ensure optimal sample quality, necropsy and sample collection were performed from ten recently deceased Kittiwakes. The birds were inspected, measured, and photographed, and their body condition score was assessed. From each bird, tracheal and cloacal swabs and a panel of tissues were collected for virological and pathological analyses, including histopathology, immunohistochemistry (IHC), RNAscope *in situ* hybridization (RNA-ISH), Maackia amurensis lectin II (MAL-II) staining, real-time reverse transcription – polymerase chain reaction (rRT-PCR), virus whole genome sequencing (WGS) and sequence analyses.

## Results

### Extent of the Outbreak

The first PCR detection of HPAI H5N1 clade 2.3.4.4b virus in Kittiwakes in Norway was recorded in May 2023 in Harstad (67°0’ N, 16°28’ E), where a smaller outbreak occurred. From May 1 to July 31, 2023, 32 of 40 dead Kittiwakes submitted for analysis at NVI tested positive for HPAI H5N1, confirming that HPAI was present in most breeding colonies in Troms and Finnmark (Figure 1).

**Figure 1.**
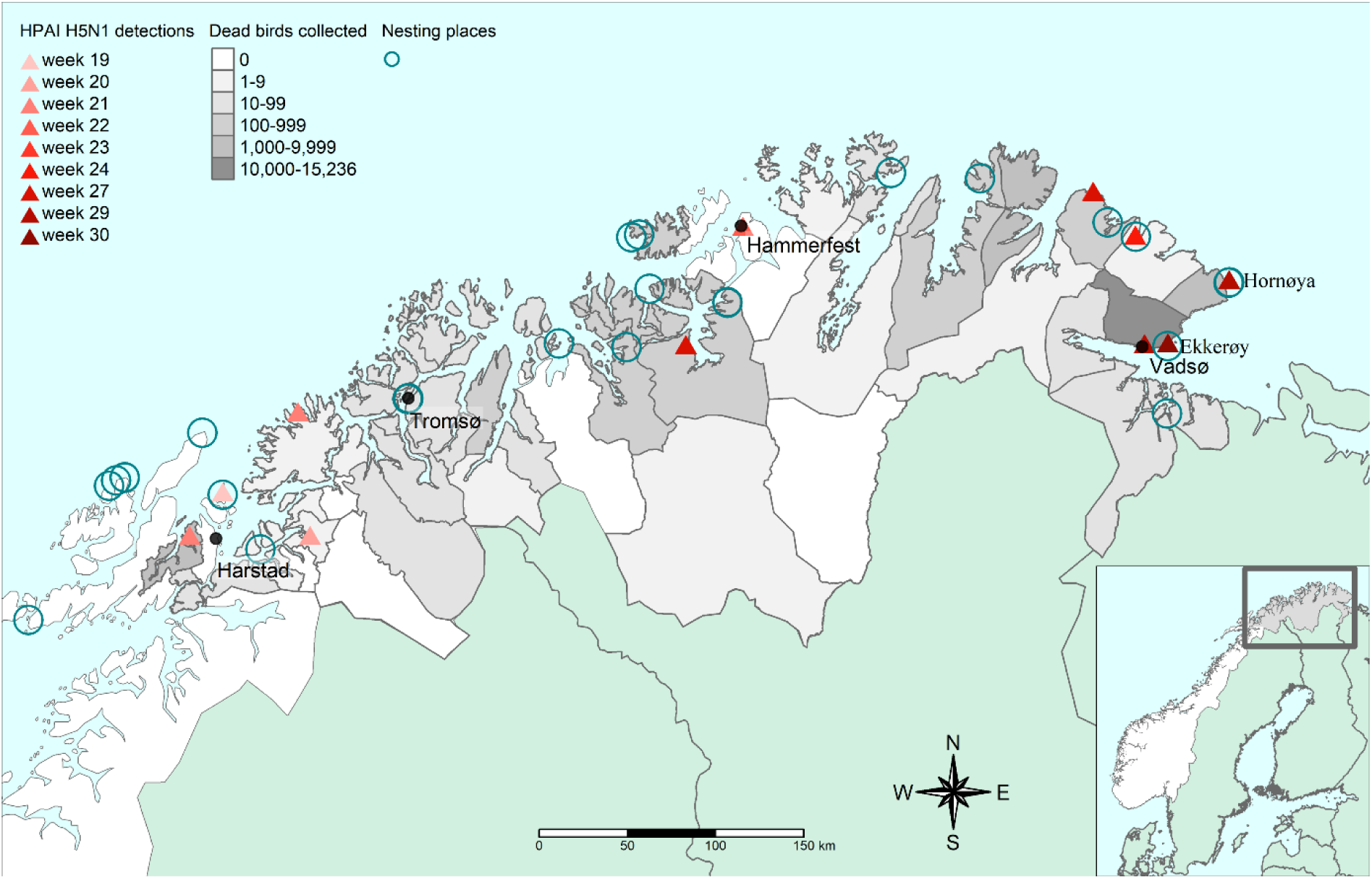
Weekly distribution of HPAIV H5N1 detections and reported mortality in Black-legged Kittiwakes (*Rissa tridactyla*), from May to September 2023, Troms and Finnmark, Norway.

Between July 28 and August 28, 2023, 24,594 wild bird carcasses, mainly Kittiwakes, were recorded in Troms and Finnmark county in Northern Norway, representing a major HPAI outbreak (Appendix 1 Table 1). Of these, 15,235 carcasses were collected in Vadsø municipality, where the Kittiwake colony at Ekkerøy is located. Large numbers of additional dead Kittiwakes were observed in areas near the colony at Ekkerøy, including coastal waters, roads, and rooftops, indicating that the true number of dead birds was even higher. For comparison, the total number of Kittiwake breeding pairs at Ekkerøy in 2023 was estimated to be 7,817 (15,634 Kittiwakes) based on colony counts (Appendix 1 Table 2).

### Clinical Characteristics

Clinically diseased Kittiwakes exhibited general lethargy and severe neurological signs, including disturbed balance, trembling, falling over, or swimming in circles. Observations included opisthotonus or head twitching, squinting eyes, and unilateral wing lameness, progressing to paralysis, paresis, recumbency, and death (Figure 2, panels A, B, <u>Video</u>).

**Figure 2.**
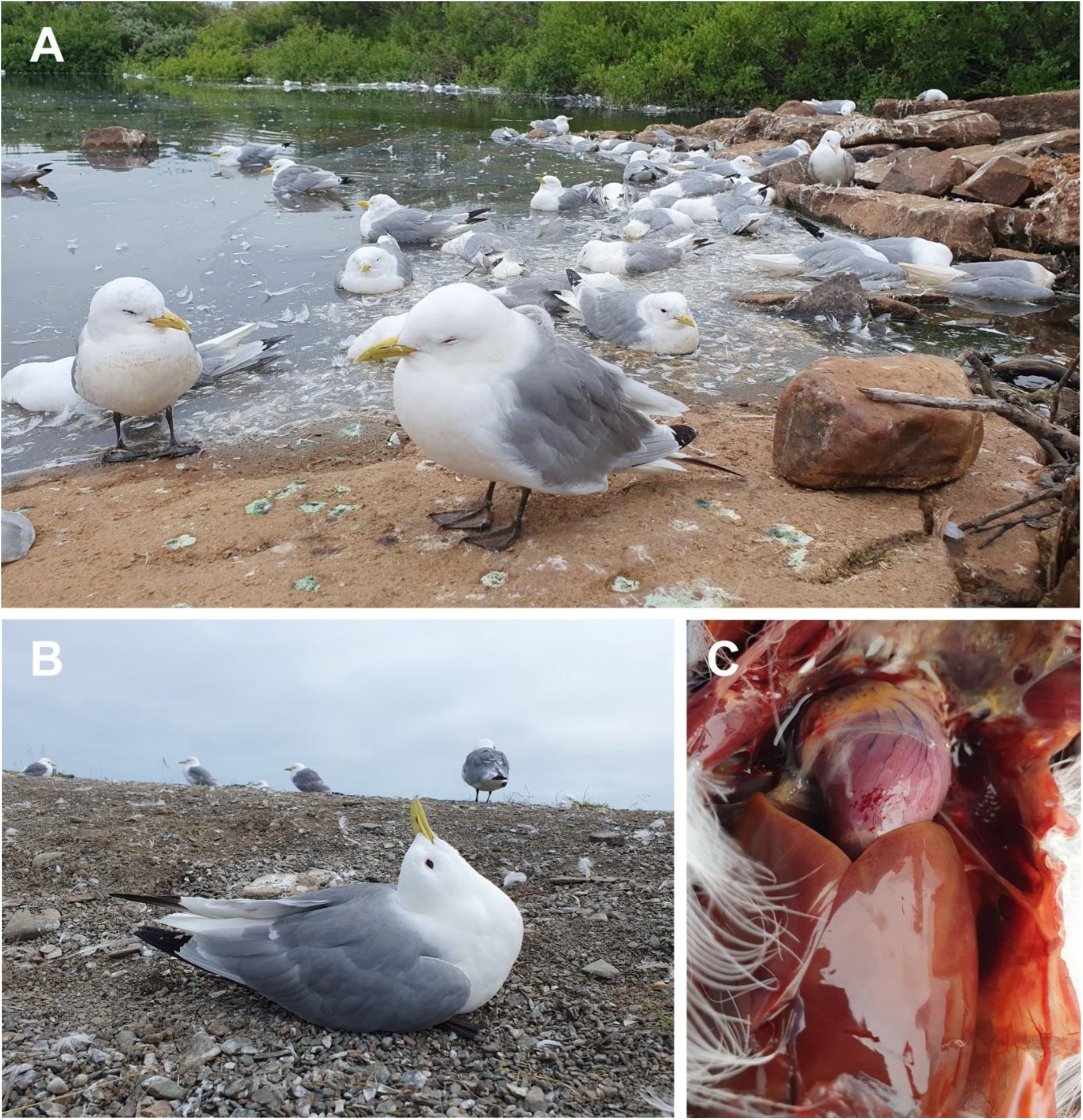
Photos taken during fieldwork at sampling point at the Storelva River near Ekkerøy, Norway, on July 26, 2023. A: Kittiwakes suffering from HPAI H5N1, several with squinting eyes and signs of apathy. In the background, dead birds and feathers floating in the water. B: Kittiwake with opisthotonus (star gazing). C: Epicardial petechiae and serous fluid in the body cavity and air sacks.

### Necropsy

The necropsied Kittiwakes, six females and four males, were adults. Two were in good condition, four were in moderate condition, and four were emaciated (Appendix 1 Table 3). Internal inspection revealed ample amounts of serous fluid in the body cavity and air sacks of all birds. Large areas of necrosis and hemorrhages in the pancreas, epicardial petechiae, and splenomegaly were observed (Figure 2, panel C).

### Histopathology

All ten birds displayed mild to moderate histopathological changes in the brain, affecting both cerebrum and cerebellum, with multifocal vascular damage, hemorrhages, vacuolization of neuropil, and multifocal necroses (Figure 3 panels A-I, Appendix 1 Table 4). This varied from single cell degeneration and necrosis of ependymal cells, neurons, and neuroglia to large areas of necrosis. Two birds had multifocal mononuclear meningitis. Apart from this, inflammatory cells were not observed. All the birds had multifocal hemorrhages and moderate to severe and multifocal-to-diffuse necroses in the pancreas (Figure 3, panels J-L, Appendix 1 Table 4). In the liver, there were mild to moderate multifocal necroses in all but one bird (Figure 3, panels M-O). No pathology was observed in the myocardium, except in one bird, which had focal subendocardial hemorrhages. No specific findings were observed in lung, spleen, proventriculus, or kidney.

**Figure. 3.**
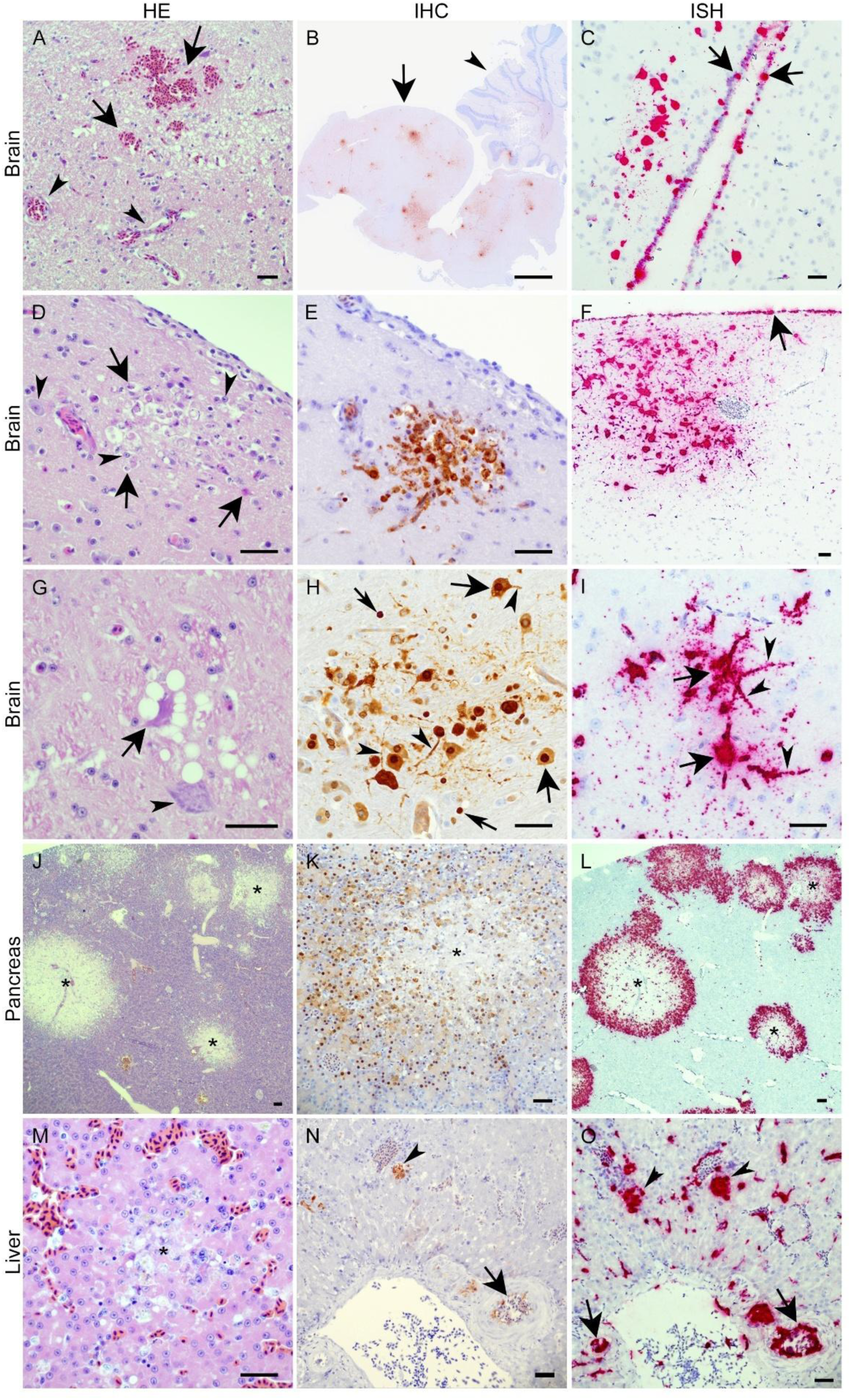
Histopathology and tissue expression of influenza A virus protein and RNA in organs from deceased Kittiwakes. Brain (A-I): Hematoxylin and eosin (HE) sections (A, D, G) demonstrated hemorrhages (A, arrows) and vacuolization of neuropil. Note intact blood vessels (A, arrowheads). Necrosis with karyorrhexis and pyknosis (D, arrows) next to viable neurons and neuroglia (D, arrowhead). Single cell necrosis with shrunken hyper-eosinophilic neuron and vacuolized neuropil (G, arrow) and viable neuron (G, arrowhead). Immunohistochemical detection of influenza A nucleoprotein (B, E, H, brown labelling) was observed multifocally in both cerebrum (B, arrow) and cerebellum (B, arrowhead). Positive labelling for influenza A nucleoprotein (E) in area of necrosis shown in serial section (D). Both neurons (H, broad arrows) and neuroglia (H, narrow arrows) were labeled, in both cell nuclei and cytoplasm. Note that the cytoplasm of both cell body and dendrites was labeled (H, arrowheads). In situ hybridization of influenza A virus RNA scope (C, F, I, red labelling) was detected multifocally in cerebellum and cerebrum in both neurons, neuroglia, and ependyma (C, arrows), meninges (F, arrow). Both cell body (I, arrows) and dendrites (I, arrowheads) of neurons were labelled. Pancreas (J-L): HE stained sections of pancreas demonstrated multifocal necroses (J, *). Labelling for Influenza A virus nucleoprotein (K) and influenza A virus RNA (I) was detected adjacent to the necrotic (*) foci in pancreas. Liver (M-O): HE stained sections of liver demonstrated areas of necrosis (M, *) with hypereosinophilic hepatocytes and karyorrhectic and pyknotic nuclei. Immunohistochemical labelling for influenza A nucleoprotein (K, brown labeling) was observed in both hepatocytes (N, arrowhead) and endothelial cells (N, arrow) of blood vessels. *In situ* hybridization demonstrated a more widespread labelling of hepatocytes (O, arrowheads), endothelial cells of blood vessels (O, arrow) and endothelial cells of hepatic sinusoids. Scale bars of A, C – I indicate 200 µm, while scale bar of B indicates 1000 µm.

### Immunohistochemical Detection of Influenza A Virus Nucleoprotein

IHC staining for influenza A virus nucleoprotein (NP) in brain tissues revealed positive labelling multifocally in and adjacent to necrotic areas, in a large number of individual neurons, glia cells, ependymal cells, and in a lower number of meningeal and vascular endothelial cells (Figure 3, panels B, E and H). Evaluation of pancreas revealed positive signal multifocally, and commonly adjacent to the necrotic foci (Figure 3, panel K). In the liver, positive signal was present in hepatocytes and endothelial cells of sinusoidal capillaries and larger blood vessels (Figure 3, panel N). No signal was detected in lung, spleen, or heart muscle.

### *In situ* detection of Influenza A Viral RNA

RNA-ISH of brain, pancreas, liver, and spleen tissues from selected birds revealed signal of various intensity, indicating the presence of viral RNA (Figure 3 panels C, F, I, L and O). The labelling was strong in cerebrum, cerebellum, and pancreas, aligning with the IHC, albeit stronger and more widespread in the vascular endothelium. Labelling of the liver varied across individuals, from sparse pinpoint signal to strong, multifocal labelling of hepatocytes and endothelial cells throughout the tissue. Spleens from two birds were examined for viral RNA, one displaying weak positive signal multifocally and the other being negative.

### Distribution of Virus Receptors

MAL-II labelled the surface of vascular endothelial cells in all examined Kittiwake organs, suggesting the presence of α-2,3-linked sialic acids (SAs) that function as AIV receptors. Labelling in the brain was sparse and confined to vascular, ependymal, and meningeal structures, but no convincing signal was observed in neurons or glial cells (Figure 4, panels A, B). Labelling was present in mucus-secreting epithelium of proventriculus and lung epithelium (Figure 4, panels C, D).

**Figure 4.**
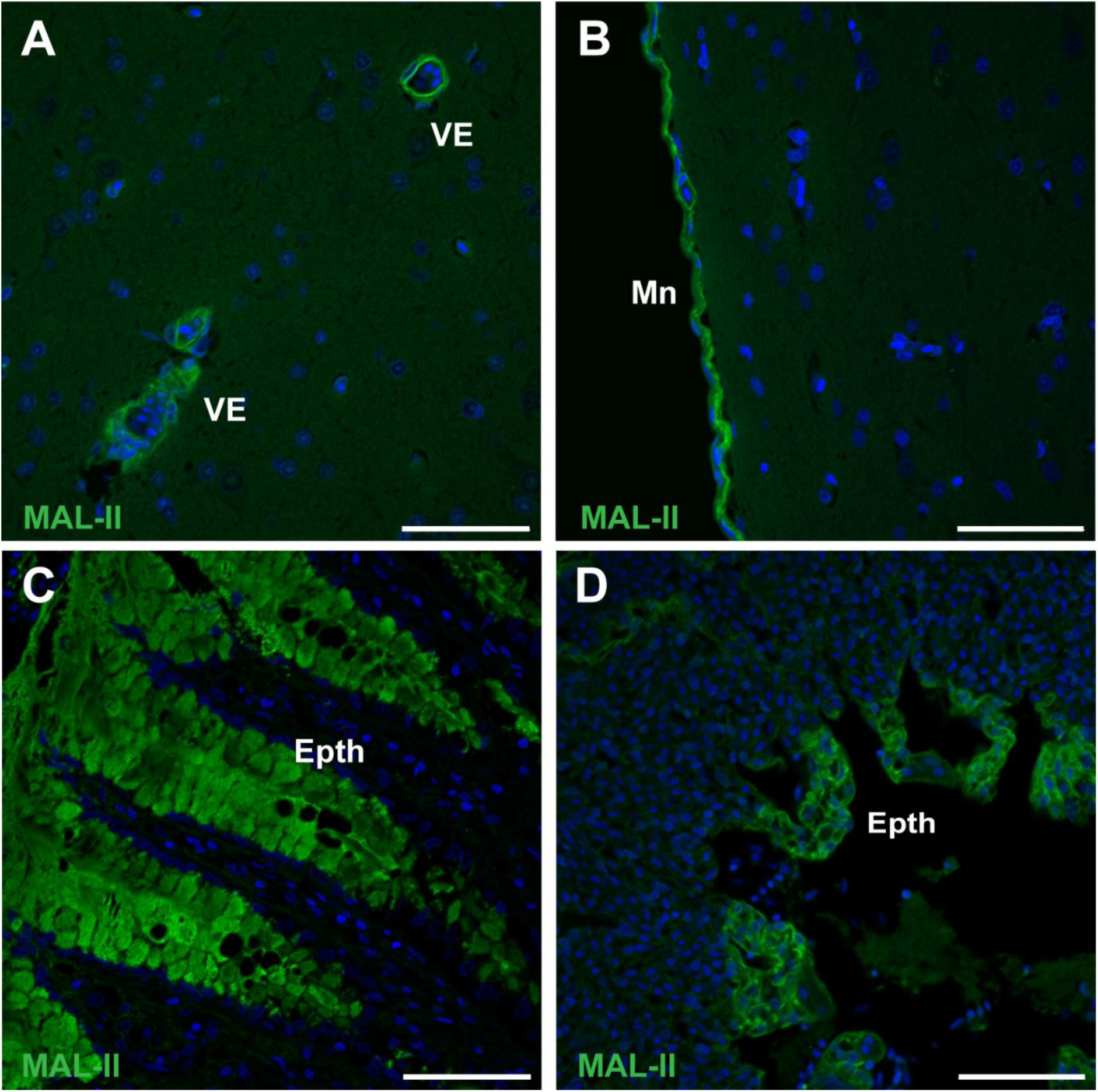
Lectin staining of presumptive avian influenza virus receptors. MAL-II (green) was used to label α-2,3-linked sialic acids in brain (A-B), lung (C), and proventriculus (D) of deceased Kittiwakes. Nuclei are labelled by Hoechst 33342 (blue). Epth = epithelium, VE = vascular endothelium, Mn = meninges. Scale bars indicate 50 µm.

### Virological Investigation

High virus levels were detected in swab and tissue samples from all individuals by rRT-PCR. The highest mean level, indicated by the quantification cycle (Cq) values, was detected in brain (Cq 15.15), followed by tracheal swabs (Cq 21.8), cloacal swabs (Cq 23.6), liver (Cq 24.8), and heart (Cq 26.0) (Figure 5).

**Figure 5.**
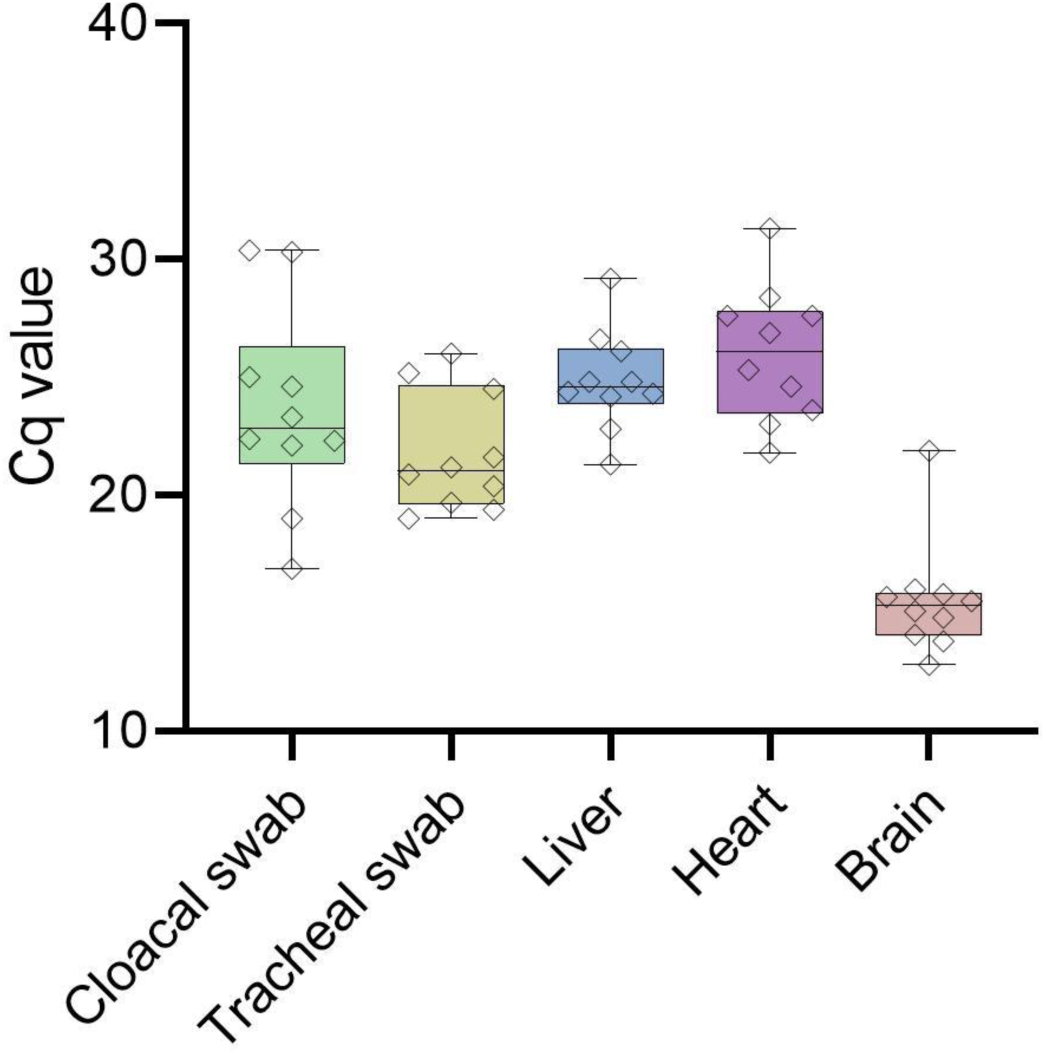
Box-and-whisker plot showing the quantification cycle (Cq) values from rRT-PCR for influenza A virus M gene performed on swab and tissue samples collected from deceased Kittiwakes (n=10), Ekkerøy, Norway, July 2023. The central line represents the median, with the whiskers indicating the range of maximum and minimum Cq values. The individual measurements are indicated by diamonds.

HPAI H5N1 clade 2.3.4.4b viruses with hemagglutinin (HA) cleavage site PLREKRRKR/GLF were confirmed in all birds, using rRT-PCR and WGS. Similarity searches in GISAID BLASTn showed that the viruses from Kittiwakes at Ekkerøy shared 99-100% identity with contemporaneous genotype BB viruses from Europe. The phylogenetic analysis showed that the viruses clustered together and were highly similar to other viruses in gulls circulating in Norway, as well as Europe during spring and summer 2023 (Figure 6, Appendix 1 Figure 2 A-G).

**Figure 6.**
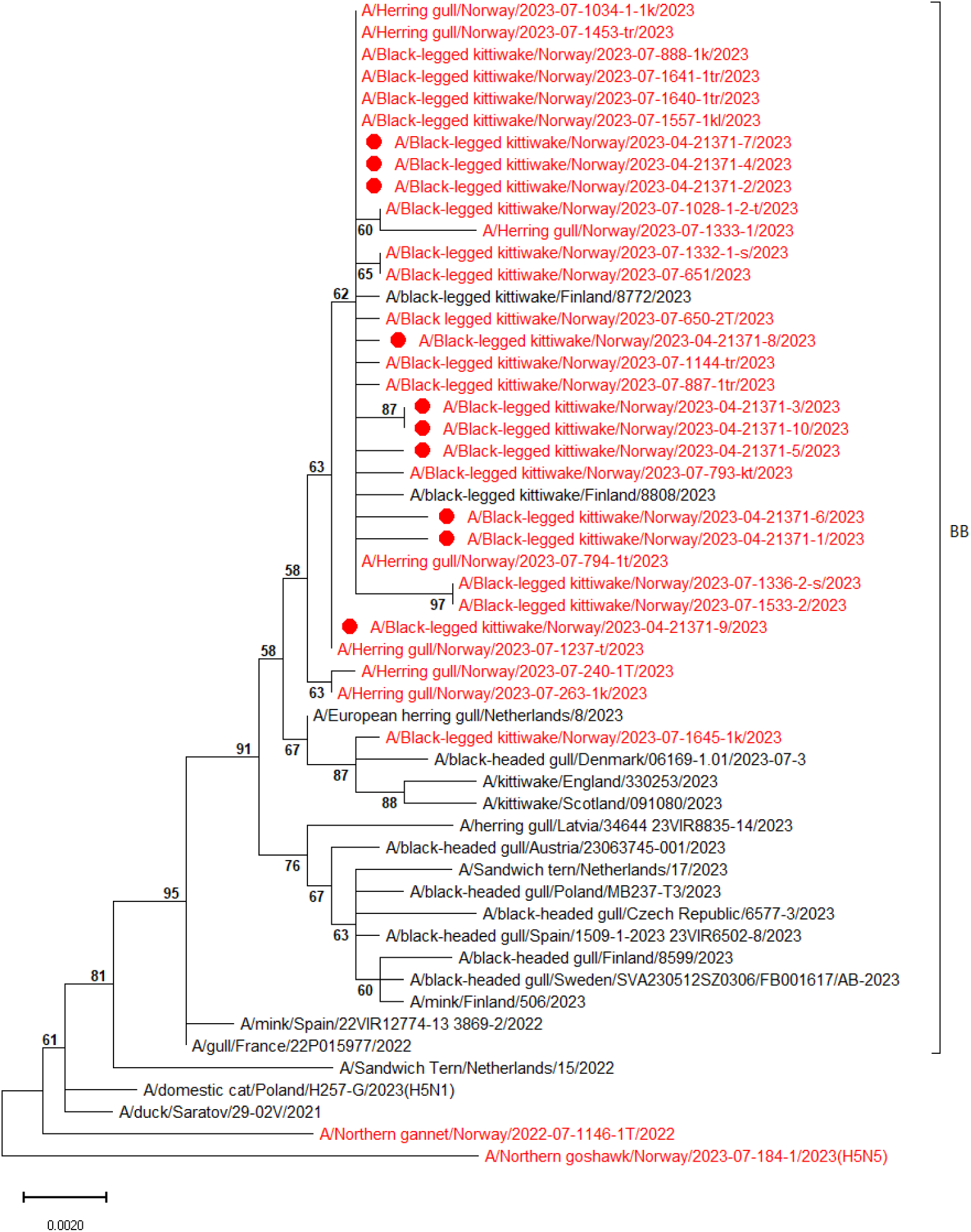
Midpoint rooted phylogenetic tree of the hemagglutinin (HA) gene from ten HPAI H5N1 viruses identified in dead Black-legged Kittiwakes from Ekkerøy, Norway, in 2023 (red dots), including contemporary genotype BB viruses from Norway (red) and Europe (black). Bootstrap values >50 are shown.

Mutation analyses showed that the viruses detected in Ekkerøy Kittiwakes remained mainly bird-adapted and had a typical profile of BB genotype, including NP-Y52N and M2-A30S (Appendix 3). Key substitutions in the HA or polymerase proteins previously associated with increased mammalian adaptation (4, 19) were not identified. No neuraminidase mutations associated with oseltamivir resistance were present.

### Management and Outbreak Response

During the outbreak, the municipality organized the removal and registration of carcasses. People handling the birds used disposable gloves, coveralls, goggles, and FFP3 masks.

Compared to Ekkerøy, the observed mortality in the neighboring Kittiwake breeding colony at Hornøya in Vardø municipality was low (observation, Tone Reiertsen). To prevent disturbance of vulnerable birds and further virus spread, movement restrictions to Hornøya and two other nature reserves were prioritized and implemented on July 27, 2023 (21). At Ekkerøy, signs providing information about the outbreak and discouraging traffic in the area were posted. No restrictions were imposed on livestock grazing in the area.

## Discussion

This study documents mass mortality in Kittiwake caused by HPAI H5N1 and enhances the understanding of the disease, pathology, and AIV receptor distribution in this species. Our estimate of the Ekkerøy breeding population size prior to the outbreak is associated with uncertainty, as it does not take into account immature individuals and “floaters” (i.e. sexually mature birds that do not breed). This implies that Ekkerøy’s total population in 2023 might have included as many as 30,000 Kittiwakes (22). Hence, with 15,235 reported dead birds and an assumption of a significant number of unreported cases, HPAI is likely to have caused a colony population reduction of at least 50%.

Seabird populations grow slowly due to their low reproduction rates and the years they spend at sea as immatures before breeding. Acute declines can severely affect future population status (23). The recovery potential depends on long-term trends in the population. Declining populations rarely recover, and the time to extinction is usually shortened (24). The HPAI outbreak may therefore have dramatic long-term impacts on the Kittiwake population in affected areas.

Clinical presentation and pathological lesions of HPAI vary by species and virus strain (25). The clinical signs of the infected Kittiwakes indicate a disease prominently affecting the central nervous system (CNS), aligning with the neurotropic nature of the virus (26–34).

A pathological finding commonly described in gulls infected with HPAI virus (HPAIV) is a sub-acute disease with lymphoplasmacytic encephalitis and areas of gliosis (28, 29, 35). This differs from our findings, where deceased Kittiwakes demonstrated widespread necrosis in the brain, pancreas, and liver, with no or limited sign of inflammation. This could suggest hematogenous spread of the HPAI H5N1 genotype BB infection with limited time for an inflammatory reaction prior to death, defining it to be a peracute, highly pathogenic, systemic, and fatal disease in Kittiwakes. One study detected considerable species variation in the severity of CNS inflammation among wild birds naturally infected with HPAI H5N1, ranging from severe in swans and Canada geese to undetectable in a Herring gull (29). Experimental inoculation of ducks and wild caught Laughing Gulls (*Larus atricilla*) with HPAI H5N1 virus produced necrosis in brain and pancreas of birds that died or were euthanized due to severe illness, corresponding to our findings in Kittiwakes (27). In birds that recovered, mild lesions such as lymphoplasmacytic perivascular encephalitis and heterophilic pancreatitis were demonstrated. As our study only included deceased birds, we do not know if any infected birds recovered from the infection, and if these then would present similar sub-acute inflammatory lesions.

Aligning with hallmark features of HPAI, the Kittiwakes showed signs of widespread systemic infection, with detection of HPAI H5N1 viral RNA in brain, liver, and heart, in addition to trachea and cloaca. By the time of death, the highest viral levels were detected in the brain, supported by low Cq values and extensive labelling by IHC and RNA-ISH. This corresponds with the histopathological lesions being most evident in this organ. Widespread histopathological lesions, in addition to extensive IHC and RNAscope labelling, indicate high virus levels also in pancreas. While we did not perform rRT-PCR on pancreatic samples, we recommend that this is included in future studies.

Virus attachment to cells is a key factor for determining viral tropism, with AIV HA preferentially binding α2,3-linked SAs (36). HPAIV replication in endothelial cells is common in chickens and swans, but less so in ducks and mammals (37). Interestingly, α2,3-linked SAs were prominently detected on vascular endothelial cells in Kittiwakes. Vascular damage with hemorrhages and endothelial expression of viral RNA and antigen were observed in brain and pancreas. This indicates that Kittiwake endothelial cells are susceptible to HPAIV, aligning with the hypothesis of hematogenous spread.

Despite the strong expression of influenza A viral RNA and NP in the brain, neurons were not labelled by MAL-II, suggesting that H5N1 targets structures in neuronal cell types that do not overlap with MAL-II binding sites (Sia-α2,3-Galβ1-GalNAc). Gangliosides are the main carriers of both α2,3– and α2,6-linked SAs in the CNS and support the attachment of different influenza A virus strains (38, 39). While comparative studies of non-infected Kittiwake tissues would be interesting, samples from healthy Kittiwakes were not feasible due to their endangered status in Norway. The entry point of the virus could not be determined in our study. Like previous observations in Herring gulls, Laughing gulls, and Ring-billed gulls (*Larus delawarensis*) (40), the mucus-secreting epithelium of Kittiwake proventriculus and lung epithelium expressed MAL-II targeted sialic acids.

The genetic similarity of the HPAI H5N1 viruses detected in our study to genotype BB viruses circulating in Europe during spring 2023, suggests introduction to breeding colonies in Northern Norway by migratory birds from Europe. The viruses in Kittiwakes were highly similar to each other and clustered with BB genotype viruses detected in other gull species in Norway, showing that the outbreak at Ekkerøy was part of a larger epizootic. We cannot exclude virus introduction to Kittiwakes during their pelagic phase in the Atlantic Ocean when they mix with other seabirds. However, we have found no data to support this, and such an introduction would have required a longer presymptomatic phase of infection. Two HPAI virus detections in Kittiwakes in Finland in July 2023 clustered among the viruses from gulls in Northern Norway, indicating same origin. Another virus from Southern Norway in July 2023 (A/Black-legged_kittiwake/Norway/2023-07-1645-1k/2023) clustered with HPAI H5N1 viruses detected in gulls in Denmark, England, and Scotland summer of 2023 and probably represents a separate introduction from the North Sea region.

Kittiwake mortality varied considerably between locations, for instance between Ekkerøy and Hornøya. A similar pattern was seen in tern colonies in the German Wadden Sea area in the summer of 2022, with some sites experiencing up to 40% mortality, and others mostly spared (41). Factors influencing mortality numbers include host susceptibility, colony size, species diversity, behavioral changes, bird movement patterns, immune status, timing, magnitude of virus introduction, environmental factors and variating recording (42–44). Understanding transmission within and between colonies requires wildlife community surveillance focusing on all interacting species.

Several factors likely contribute to the severity of the HPAI outbreaks in Kittiwakes. Due to the pelagic lifestyle of Kittiwakes, pre-existing immunity to H5 is assumed to have been low prior to the epizootic in 2023 (16, 19). Virus properties, specifically the genetic constellation of the BB genotype also likely enhanced virus replication in gulls (45). Kittiwakes nest closely together, facilitating virus transmission. The outbreak at Ekkerøy coincided with a high number of susceptible chicks, and adults with a reduced body condition towards the end of the breeding period that could have lowered their resistance and resilience to infection. At the Storelva River near Ekkerøy, numerous infected carcasses, feathers, and feces combined with low water levels likely promoted virus spread through freshwater. Environmental samples could have provided more information and should be obtained in future outbreaks.

The viruses detected in Kittiwakes at Ekkerøy were avian adapted with mutation profiles similar to other BB genotype viruses, thus not indicating elevated risk for mammal adaptation. However, a few mutations that could increase the zoonotic risk were present. NP-Y52N associated with evasion of human BTN3A3 restriction was identified (46).

Mitigation measures can be implemented to reduce HPAI spread and protect vulnerable wild bird populations. One measure is removal of carcasses, but this must be balanced against the risk of disturbing the remaining birds in the colony. The effect of carcass removal is difficult to measure. A study of HPAI-affected Sandwich tern (*Thalasseus sandvicensis*) colonies found that carcass removal reduced adult mortality by an average of 15% (47). To keep the infection pressure in the environment low and reduce spillover to other birds and mammals, carcass removal should start early in an outbreak and be performed by trained personnel using personal protective equipment. The high number of carcasses at Ekkerøy was challenging to handle due to limited staff, underlining the need for joint effort in extreme situations. Management of HPAI mass mortalities in wildlife requires a multisectoral One Health approach, with clear organizational roles.

Our study does not determine to what extent birds survived infection and may have gained immunity. Studies that can provide such data may be of great value for projecting the vulnerability of this and other Kittiwake colonies to future A/H5Nx HPAI outbreaks.

## Conclusion

We recommend further monitoring of the long-term impact of HPAI on the Kittiwake populations. HPAI outbreaks in seabird colonies need awareness and must be mitigated using a multi-sectorial One Health approach to protect endangered wildlife species, mammals and public health.

## Acknowledgements

We gratefully acknowledge all data contributors, i.e., the authors and their originating laboratories responsible for obtaining the specimens, and their submitting laboratories for generating the genetic sequence and metadata and sharing via the GISAID Initiative, on which this research is based. We would like to thank Knut Sverre Horn for photographing the colony at Ekkerøy and Christer Michalsen and Hans Ivar Hortman, both from the Norwegian Nature Inspectorate, for making this possible. We thank Jennifer Stien for help with counts from the photos. We also thank the County Governor of Troms and Finnmark for sharing the mortality data and the Norwegian Food Safety Authority and collaborating organizations for submitting samples for the passive surveillance program for avian influenza. We are grateful to all laboratory personnel at the NVI who contributed to analyzing the samples and thank Bjørnar Ytrehus (NVI) for performing body condition score assessments.

