## Appendix 1 for "Highly Pathogenic Avian Influenza A(H5N1) Caused Mass Death among Black-legged Kittiwakes (*Rissa tridactyla*) in Norway, 2023"

#### Materials and methods

*Organizational structure*

The Norwegian Food Safety Authority (NFSA) is the administrative authority for the health and welfare of wild and domestic animals in Norway, and responsible for the surveillance programme for HPAI in wild birds. Wildlife managers, ornithologist and citizens notify the NFSA when HPAI is suspected. The NFSA or their collaborators collect swabs from the trachea and cloaca of the birds and ship the samples to the Norwegian Veterinary Institute (NVI) for analysis. NVI is the national reference laboratory for avian influenza and is responsible for planning laboratory investigation and reporting components of the programme. In addition, NVI conducts risk assessments and provides advice on disease prevention and management in case of outbreaks in animals. The Norwegian Institute of Public Health (NIPH) is responsible for influenza in humans, gives advice on public health aspects and is the national reference laboratory for human influenza as well as the WHO National Influenza Centre.

*Epidemiological analysis*

To provide an epidemiological description of the outbreak in northern Norway, data on HPAI virus (HPAIV) PCR detections in Black-legged Kittiwakes (*Rissa tridactyla*) were obtained from the laboratory reporting system at the NVI. Data were analysed and maps created using R version 4.3.0 (1). Data of locations of Kittiwake nests were obtained from the Species Observations System (2).

*Population estimates*

During the outbreak, the colony was documented through photography (Figure 1), and the total number of apparently occupied nests was recorded to estimate the population of Kittiwake breeding pairs present in 2023 (Table 2). Nests containing chicks or adults or both were counted as occupied nest. In addition, nests with dead chicks or adults or empty nests with current years' nesting material and traces of fresh faeces were counted as occupied. Nests with an uncertain status were included in the final estimate.

*Necropsy*

During fieldwork in July 2023, necropsy and sample collection were performed from ten recently deceased Kittiwakes. The body condition was determined by scoring the pectoral muscle based on photographs in accordance with the description in Franeker J.A. van & C.J. Camphuysen 2007 (3). Swabs (COPAN, eSwab) from trachea and cloaca, and tissue from brain, liver and heart (5 x 5 x 5 mm) were collected separately into tubes containing UTM-medium for PCR analyses at NVI. For histopathology, immunohistochemistry, RNAscope- and receptor analyses whole/half brain and approximately 1 x 1 x 0.5 cm large tissue samples were collected from heart, lung, liver, kidney, spleen, pancreas and proventriculus and placed in 10% buffered formalin.

*Histopathology, immunohistochemistry and RNAscope* in situ *hybridization*

Tissue samples fixed in 10% buffered formalin, were subsequently dehydrated in ethanol, equilibrated in xylene and embedded in paraffin. For histology, 2-3 µm thick sections were stained with hematoxylin and eosin (HE) and examined by a trained pathologist. In addition, selected tissue sections were subjected to immunohistochemistry; labelled with antibodies against influenza A nucleoprotein (NP) (Anti Influenza A NP, mouse monoclonal IgG1, produced on virus particles from H5N2 strain, Statens Serum Institut). 2-3 µm tissue sections were mounted onto poly-lysine-coated slides, heated to 60 ºC for at least 30 minutes and deparaffinized. The slides were blocked for endogen peroxidase for ten minutes followed by enzyme treatment applying 2 µl proteinase K recombinant PCR Grade in 4 ml TRIS-buffer applied to the sections for 30 minutes at 37 ºC. Subsequent step included blocking with 5 % BSA in TRIS buffer solution for 20 minutes. The primary anti-influenza A NP antibody was diluted 1:2000 for one hour at room temperature, followed by application of the secondary antibody EnVision HRP anti-mouse for 30 minutes at room temperature. Substrate used for visualization was Romulin AEC chromogen, mixing 16 µl solution A, B and C in 2.5 ml Romulin buffer, 15 minutes at room temperature.

Paraffin-embedded tissue sections were used to detect viral sequences through the RNAscope™ *in situ* hybridization (ISH) technique (4). RNAscope ISH is an innovative method that allows the visualization of individual RNA molecules in cytological and histological samples at the single-cell level. In our study, we employed a probe that targets the matrix gene of the Avian Influenza A Virus (AIAV), which is conserved across all influenza A viruses. This probe was obtained from ACD Bio (RNAscope™ Probe - V-InfluenzaA-H5N8-M2M1-C1, Cat. Nr. 1048221-C1), and the slides were stained using the RNAscope™ 2.5 HD Detection Reagents-RED assay (ACD Bio, Cat. Nr. 322360). For the negative control, we used a probe that targets the dihydrodipicolinate reductase (dapB) gene from *Bacillus subtilis* (Cat No. 310043). For the positive control, RNAscope® Probe - Gg-PPIB - *Gallus gallus* peptidylprolyl isomerase B (cyclophilin B) (PPIB) mRNA (Cat No. 453371) was utilized, which serves as a housekeeping gene.

*MAL-II lectin staining*

To map the distribution of H5N1 virus receptors, the binding pattern of the lectin MAL-II was assessed (5). Avian influenza A viruses bind Siaα2–3Galβ1-terminated glycans, with further diversity in binding specificity depending on underlying glycan types and modifications (6). The MAL-II lectin specifically binds Siaα2–3Galβ1-3GalNAcs (7). Tissue sections from brain, lung and proventriculus (4 µm) were deparaffinized and incubated MAL-II (CliniSciences, France; 40 µg/mL in PBS) for 60 min at room temperature, washed (PBS, 5 min x 2), counterstained with Hoechst 33342 (2 µg/mL, Thermo Fisher Scientific), and mounted in ProLong Glass Antifade mountant (Thermo Fisher Scientific). Confocal images were obtained using a Zeiss Axio LSM710 equipped with a Plan-Apochromat 40x/1.3 NA oil objective and Zen 2.3SP1 software.

*Real-time RT-PCR and whole genome sequencing*

Real-time RT-PCR (rRT-PCR) for influenza A virus was performed on the collected cloacal and tracheal swabs, as well as liver, heart, and brain tissue samples. The swabs and tissue samples were pretreated with MagNA Pure External Lysis Buffer (Roche, 06374913001). Approximately 20 mg of tissue was homogenized before total RNA/DNA was extracted from all samples using MagNA Pure 96 automated extractor (Roche) with the MagNA Pure 96 DNA and viral NA LV Kit (Roche, 6374891001). Detection of Influenza A virus was set up using an rRT-PCR assay targeting the conserved M gene (8), in accordance with the Manual of standards for diagnostic tests and vaccines (9). Subtyping was performed with a specific assay for the HPAI H5 clade 2.3.4.4b (10, 11) together with HA and NA subtype specific rRT-PCR-assays (12, 13). The rRT-PCR for detection and subtyping of Influenza A virus was carried out with OneStep RT-PCR Kit (Qiagen, 210212) and Brilliant III Ultra-Fast QRT-PCR Master Mix (Agilent Technologies, 600884). All analyses were run on AriaMX instruments by Agilent.

Nucleic acids extracted from the brain samples underwent cDNA synthesis, as described by Zhou et al., 2009 and were subjected to whole genome sequencing (14). Briefly, extracted nucleic acids (same as for PCR) were employed as template for cDNA synthesis and subsequent PCR using degenerate primers amplifying all 8 segments of the Influenza A virus (Superscript III Platinum one-step quantitative RT-PCR system (Thermo Fisher Scientific, Waltham, MA, USA)). The PCR products were analyzed and quantified on a TapeStation 4200 (Agilent Technologies, Santa Clara, CA, USA). DNA libraries were prepared using Illumina DNA Prep (Illumina, San Diego, CA, USA) and sequenced using Illumina technology (MiSeq). Consensus sequences were made by reference-based mapping towards A/Black_legged_kittiwake/Norway/2023-07-650-2T/2023 (EPI_ISL_18455264) in InsaFlu (15).

*Sequence analyses*

Genetic characterization of the detected viruses was performed, and BLAST (16) in GISAID (17) was used for similarity searches. Phylogenetic analysis compared the nucleotide sequences of viruses from the ten necropsied Kittiwakes with AIV sequences from Norway (March-July 2023) and selected European sequences (2020-2023) from GISAID EpiFlu. An overview of all sequences included in the analyses is given in the Appendix 2. For each segment, the full-length coding nucleotide sequences were aligned by Clustal W and phylogenetic trees were created using the Maximum likelihood algorithm and the Tamura Nei model in MEGA 10.1.8 (18, 19). We used bootstrap values of 1000 replicates to assess the nodal support. Mutation analyses were performed to search for mutations or motifs previously associated with mammalian viral adaptation or antiviral resistance, using FluMutGUI 3.1.1; FluMut 0.6.3; FluMutDB 6.3, released on 2024-09-12 (https://github.com/izsvenezie-virology/FluMut).

### Results

Table 1. Number of dead birds reported by the municipalities in Troms and Finnmark County, Norway, between July 28 and August 28, 2023, due to the HPAI H5N1 outbreak in Kittiwakes (*Rissa tridactyla*) in Northern Norway (20). An empty cell means that no data was reported.

| Municipality | July 28 | August 1 | August 4 | Week 32 | Week 33 | Week 34 | Total |
|---|---|---|---|---|---|---|---|
| Alta | 140 | 100 | | 30 | 0 | | 270 |
| Balsfjord | 8 | 2 | | 3 | | | 13 |
| Bardu | | | | | | | 0 |
| Berlevåg | 358 | | 23 | 3 | | | 384 |
| Båtsfjord | | 9 | | | | | 9 |
| Dyrøy | 0 | 0 | 2 | 5 | 0 | 0 | 7 |
| Gamvik | | 600 | | 400 | 500 | | 1500 |
| Gratangen | 0 | 2 | 0 | 0 | 0 | | 2 |
| Hammerfest | 2000 | 229 | 46 | 67 | 16 | 14 | 2372 |
| Harstad | | 5 | 2 | 8 | | | 15 |
| Hasvik | 400 | 100 | | 330 | 0 | | 830 |
| Ibestad | 6 | | 3 | 0 | | | 9 |
| Karasjok | | 0 | | | | | 0 |
| Karlsøy | 10 | 13 | 0 | 0 | 0 | 0 | 23 |
| Kautokeino | 1 | 0 | 0 | | | | 1 |
| Kvæfjord | 279 | 680 | 97 | 251 | 42 | 20 | 1369 |
| Kvænangen | 25 | 150 | 100 | 0 | 40 | | 315 |
| Kåfjord | 50 | 5 | | 0 | | | 55 |
| Lavangen | 0 | 0 | 0 | 0 | | | 0 |
| Lebesby | 500 | 40 | 0 | 8 | 0 | | 548 |
| Loppa | 100 | 81 | | 0 | | | 181 |

| | | | | | | | |
|---|---|---|---|---|---|---|---|
| Lyngen | 143 | 21 | 0 | 2 | 0 | 0 | 166 |
| Målselv | 4 | | 8 | 0 | | | 12 |
| Måsøy | | 2 | | | | | 2 |
| Nesseby | 40 | | | | | | 40 |
| Nordkapp | | 60 | | | | | 60 |
| Nordreisa | 7 | 0 | 0 | 0 | 13 | | 20 |
| Porsanger | 2 | | | | | | 2 |
| Salangen | 0 | 5 | 0 | 0 | 0 | 0 | 5 |
| Senja | 0 | 0 | 3 | 0 | 0 | 0 | 3 |
| Skjervøy | 12 | 6 | 0 | 5 | | | 23 |
| Storfjord | 5 | 1 | 2 | 0 | 0 | 0 | 8 |
| Sørreisa | 0 | 0 | 0 | 0 | | | 0 |
| Sør-Varanger | | 75 | | | | | 75 |
| Tana | 0 | 0 | 0 | 0 | 1 | | 1 |
| Tjeldsund | 2 | 4 | 1 | 4 | | | 11 |
| Tromsø | 7 | 6 | 2 | 6 | 1 | 6 | 28 |
| Vadsø | 12593 | 1543 | 891 | 208 | | | 15235 |
| Vardø | 1000 | | | | | | 1000 |
| Total | 17692 | 3739 | 1180 | 1330 | 613 | 40 | 24594 |

Table 2. Counts of apparently occupied nests during the outbreak, used for estimation of the number of breeding pairs of Black-legged Kittiwakes at the Ekkerøy colony, Norway, in 2023.

| Category | Number of nests |
|---|---|
| At least one adult and one chick on nest | 335 |
| At least one living chick but no adult(s) on nest | 669 |
| One or more adults and no chick(s) on nest | 4976 |
| Nest used in 2023, but no birds present | 1423 |
| Dead chick(s) on nest | 414 |
| Uncertainty of occupied nest | 111 |
| Total apparently occupied nests (included uncertain nests) | 7928 |
| Total apparently occupied nests (without uncertain nests) | 7817 |

Table 3. Individual data and Pectoral Muscle Condition Score (PMCS) of ten Black-legged Kittiwakes that died from HPAI H5N1 and were necropsied at Storelva, Norway, in July 2023.

| Bird no. | 1 | 2 | 3 | 4 | 5 | 6 | 7 | 8 | 9 | 10 |
|---|---|---|---|---|---|---|---|---|---|---|
| Weight (g) | 312 | 378 | 310 | 312 | 321 | 308 | 374 | 287 | 266 | 294 |
| Sex* | F | M | M | F | F | F | M | M | F | F |
| Head (cm) | 9.1 | 9.6 | - | - | 9.0 | 8.8 | 9.5 | 9.0 | 8.4 | 8.9 |
| PMCS** | 3 | 3 | 2 | 1 | 2 | 2 | 1 | 1 | 1 | 2 |

*F: Female, M: Male

**1: emaciated, 2: moderate condition, 3: good condition

Table 4. Score of histopathological examination: Hematoxylin and eosin (HE)-stained slides of organs from ten Black-legged Kittiwakes that died from HPAI H5N1 at Ekkerøy, Norway, in 2023, were examined and scored semi-quantitatively based on histopathological observations. The severity of changes was noted as: none (0), mild (1), moderate (2) and severe (3). Missing organ was noted as: (-).

| Bird number | 1 | 2 | 3 | 4 | 5 | 6 | 7 | 8 | 9 | 10 |
|---|---|---|---|---|---|---|---|---|---|---|
| Brain | 1 | 1 | 2 | 2 | 2 | 2 | 2 | 2 | 2 | 2 |
| Pancreas | 2 | 2 | 2 | 2 | 2 | 2 | 3 | 2 | 3 | 2 |
| Liver | 1 | 2 | 1 | 2 | 1 | 0 | 1 | 1 | 2 | 1 |
| Spleen | - | 0 | 0 | 0 | 0 | 0 | 0 | 0 | 0 | 0 |
| Heart | 0 | 0 | 1 | 0 | 0 | 0 | 0 | 0 | 0 | 0 |
| Kidney | 0 | 0 | 0 | 0 | 0 | 0 | 0 | 0 | 0 | 0 |
| Lung | 0 | 0 | 0 | 0 | 0 | 0 | 0 | 0 | 0 | 0 |
| Proventriculus | 0 | 0 | 0 | 0 | 0 | 0 | 0 | 0 | 0 | - |

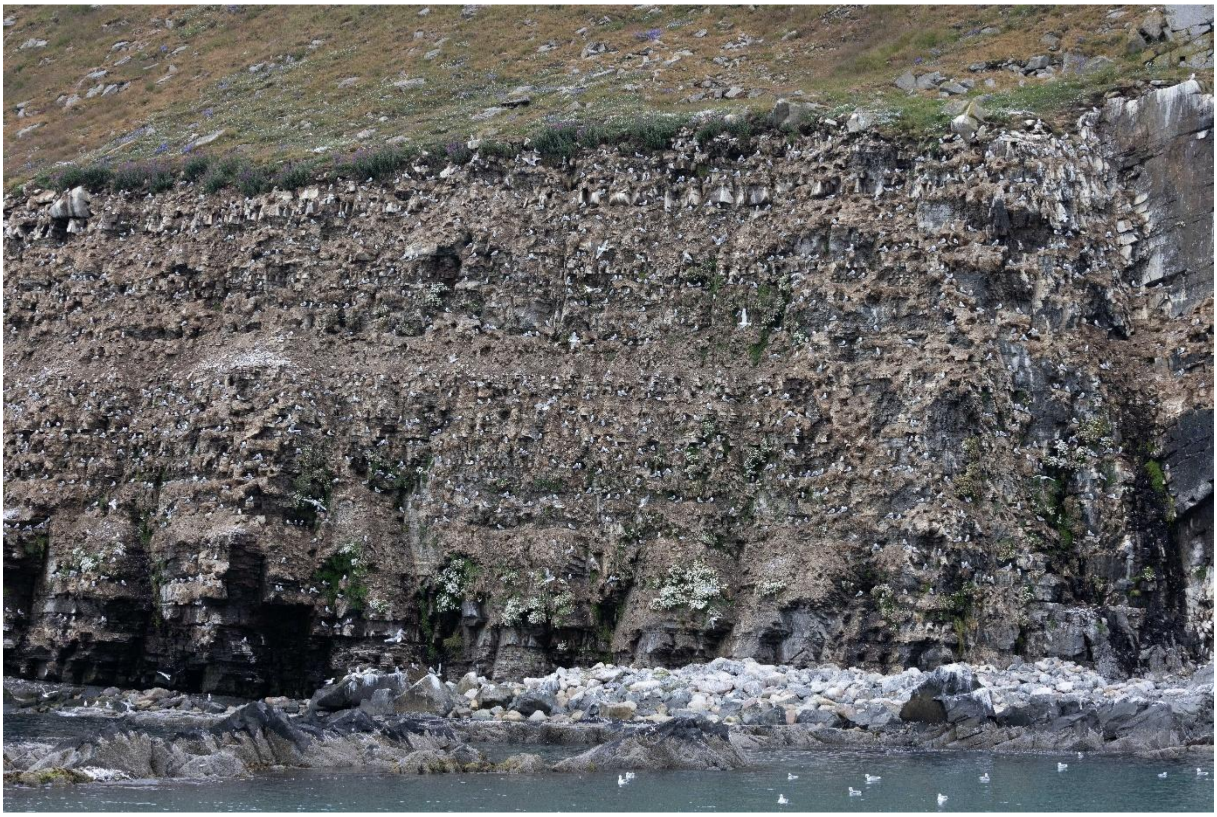

Figure 1. A photo segment of the Kittiwake cliff on Ekkerøy from July 2023, showcasing the image resolution. Apparently occupied nests were counted from several such pictures to estimate the size of the breeding population in 2023. Photo: Knut Sverre Horn.

A)

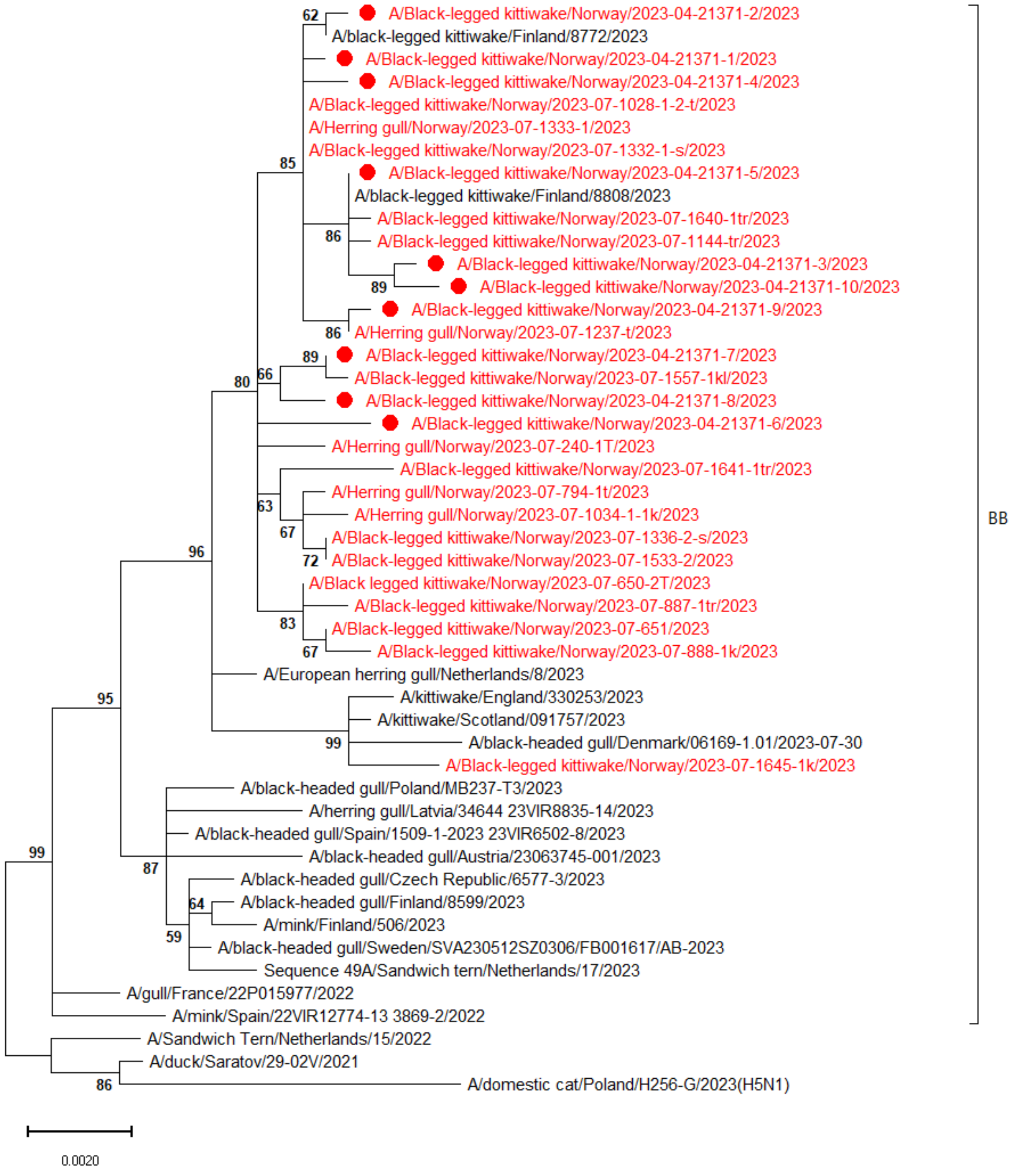

62
A/Black-legged kittiwake/Norway/2023-04-21371-2/2023
A/black-legged kittiwake/Finland/8772/2023
A/Black-legged kittiwake/Norway/2023-04-21371-1/2023
A/Black-legged kittiwake/Norway/2023-04-21371-4/2023
A/Black-legged kittiwake/Norway/2023-07-1028-1-2-t/2023
A/Herring gull/Norway/2023-07-1333-1/2023
A/Black-legged kittiwake/Norway/2023-07-1332-1-s/2023
85
A/Black-legged kittiwake/Norway/2023-04-21371-5/2023
A/black-legged kittiwake/Finland/8808/2023
A/Black-legged kittiwake/Norway/2023-07-1640-1tr/2023
86
A/Black-legged kittiwake/Norway/2023-07-1144-tr/2023
A/Black-legged kittiwake/Norway/2023-04-21371-3/2023
89
A/Black-legged kittiwake/Norway/2023-04-21371-10/2023
A/Black-legged kittiwake/Norway/2023-04-21371-9/2023
86
A/Herring gull/Norway/2023-07-1237-t/2023
89
A/Black-legged kittiwake/Norway/2023-04-21371-7/2023
80
66
A/Black-legged kittiwake/Norway/2023-07-1557-1kl/2023
A/Black-legged kittiwake/Norway/2023-04-21371-8/2023
A/Black-legged kittiwake/Norway/2023-04-21371-6/2023
A/Herring gull/Norway/2023-07-240-1T/2023
A/Black-legged kittiwake/Norway/2023-07-1641-1tr/2023
A/Herring gull/Norway/2023-07-794-1t/2023
63
A/Herring gull/Norway/2023-07-1034-1-1k/2023
67
A/Black-legged kittiwake/Norway/2023-07-1336-2-s/2023
96
72
A/Black-legged kittiwake/Norway/2023-07-1533-2/2023
A/Black legged kittiwake/Norway/2023-07-650-2T/2023
A/Black-legged kittiwake/Norway/2023-07-887-1tr/2023
83
A/Black-legged kittiwake/Norway/2023-07-651/2023
67
A/Black-legged kittiwake/Norway/2023-07-888-1k/2023
A/European herring gull/Netherlands/8/2023
95
A/kittiwake/England/330253/2023
A/kittiwake/Scotland/091757/2023
99
A/black-headed gull/Denmark/06169-1.01/2023-07-30
A/Black-legged kittiwake/Norway/2023-07-1645-1k/2023
A/black-headed gull/Poland/MB237-T3/2023
A/herring gull/Latvia/34644 23VIR8835-14/2023
A/black-headed gull/Spain/1509-1-2023 23VIR6502-8/2023
99
A/black-headed gull/Austria/23063745-001/2023
87
A/black-headed gull/Czech Republic/6577-3/2023
64
A/black-headed gull/Finland/8599/2023
A/mink/Finland/506/2023
59
A/black-headed gull/Sweden/SVA230512SZ0306/FB001617/AB-2023
Sequence 49A/Sandwich tern/Netherlands/17/2023
A/gull/France/22P015977/2022
A/mink/Spain/22VIR12774-13 3869-2/2022
A/Sandwich Tern/Netherlands/15/2022
A/duck/Saratov/29-02V/2021
86
A/domestic cat/Poland/H256-G/2023(H5N1)
BB
0.0020

B)

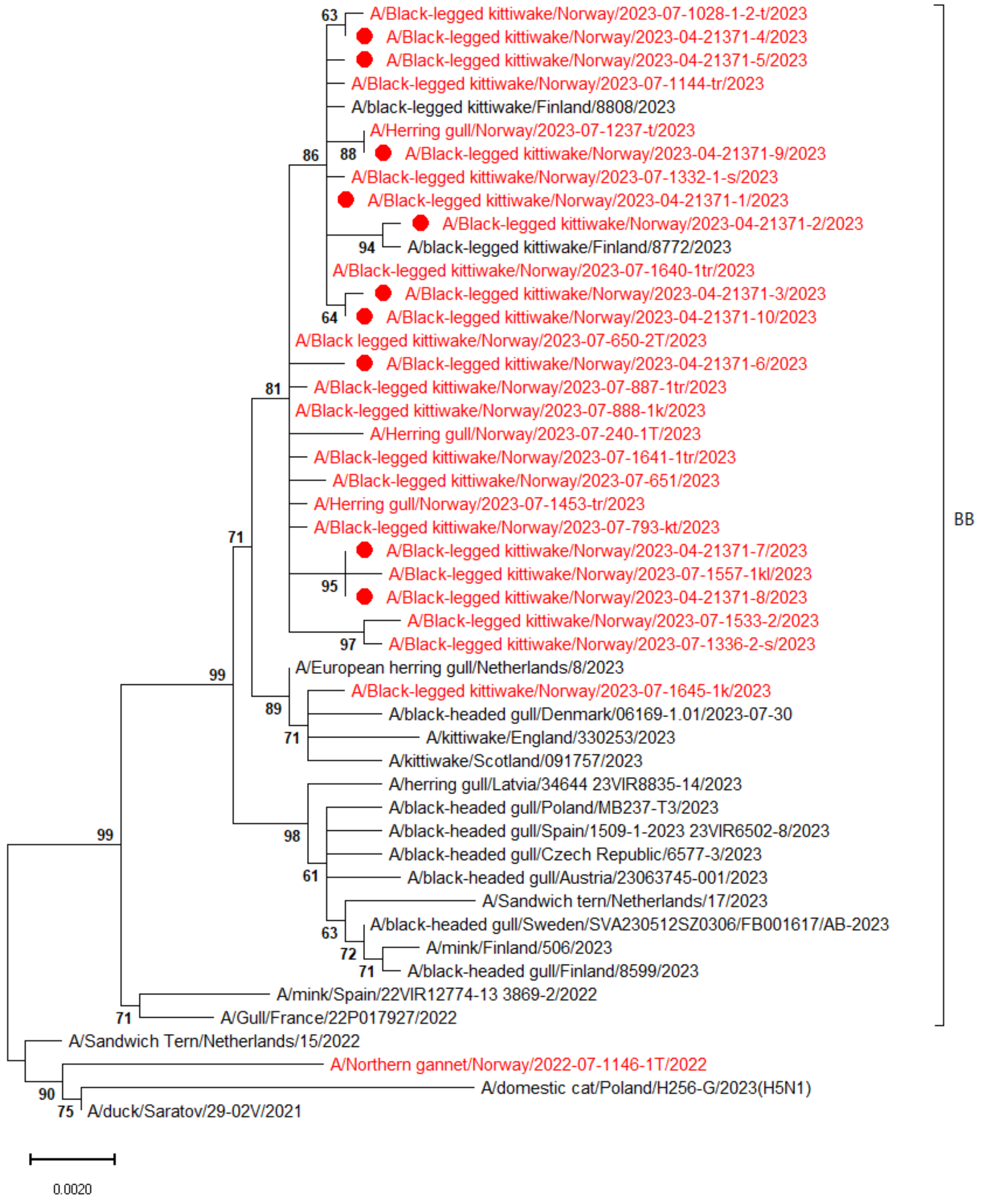
A/Black-legged kittiwake/Norway/2023-07-1028-1-2-t/2023
A/Black-legged kittiwake/Norway/2023-04-21371-4/2023
A/Black-legged kittiwake/Norway/2023-04-21371-5/2023
A/Black-legged kittiwake/Norway/2023-07-1144-tr/2023
A/black-legged kittiwake/Finland/8808/2023
A/Herring gull/Norway/2023-07-1237-t/2023
A/Black-legged kittiwake/Norway/2023-04-21371-9/2023
A/Black-legged kittiwake/Norway/2023-07-1332-1-s/2023
A/Black-legged kittiwake/Norway/2023-04-21371-1/2023
A/Black-legged kittiwake/Norway/2023-04-21371-2/2023
A/black-legged kittiwake/Finland/8772/2023
A/Black-legged kittiwake/Norway/2023-07-1640-1tr/2023
A/Black-legged kittiwake/Norway/2023-04-21371-3/2023
A/Black-legged kittiwake/Norway/2023-04-21371-10/2023
A/Black legged kittiwake/Norway/2023-07-650-2T/2023
A/Black-legged kittiwake/Norway/2023-04-21371-6/2023
A/Black-legged kittiwake/Norway/2023-07-887-1tr/2023
A/Black-legged kittiwake/Norway/2023-07-888-1k/2023
A/Herring gull/Norway/2023-07-240-1T/2023
A/Black-legged kittiwake/Norway/2023-07-1641-1tr/2023
A/Black-legged kittiwake/Norway/2023-07-651/2023
A/Herring gull/Norway/2023-07-1453-tr/2023
A/Black-legged kittiwake/Norway/2023-07-793-kt/2023
A/Black-legged kittiwake/Norway/2023-04-21371-7/2023
A/Black-legged kittiwake/Norway/2023-07-1557-1kl/2023
A/Black-legged kittiwake/Norway/2023-04-21371-8/2023
A/Black-legged kittiwake/Norway/2023-07-1533-2/2023
A/Black-legged kittiwake/Norway/2023-07-1336-2-s/2023
A/European herring gull/Netherlands/8/2023
A/Black-legged kittiwake/Norway/2023-07-1645-1k/2023
A/black-headed gull/Denmark/06169-1.01/2023-07-30
A/kittiwake/England/330253/2023
A/kittiwake/Scotland/091757/2023
A/herring gull/Latvia/34644 23VIR8835-14/2023
A/black-headed gull/Poland/MB237-T3/2023
A/black-headed gull/Spain/1509-1-2023 23VIR6502-8/2023
A/black-headed gull/Czech Republic/6577-3/2023
A/black-headed gull/Austria/23063745-001/2023
A/Sandwich tern/Netherlands/17/2023
A/black-headed gull/Sweden/SVA230512SZ0306/FB001617/AB-2023
A/mink/Finland/506/2023
A/black-headed gull/Finland/8599/2023
A/mink/Spain/22VIR12774-13 3869-2/2022
A/Gull/France/22P017927/2022
A/Sandwich Tern/Netherlands/15/2022
A/Northern gannet/Norway/2022-07-1146-1T/2022
A/domestic cat/Poland/H256-G/2023(H5N1)
A/duck/Saratov/29-02V/2021
63
86
88
94
64
81
71
95
97
99
89
71
98
61
63
72
71
99
71
90
75
BB
0.0020

C)

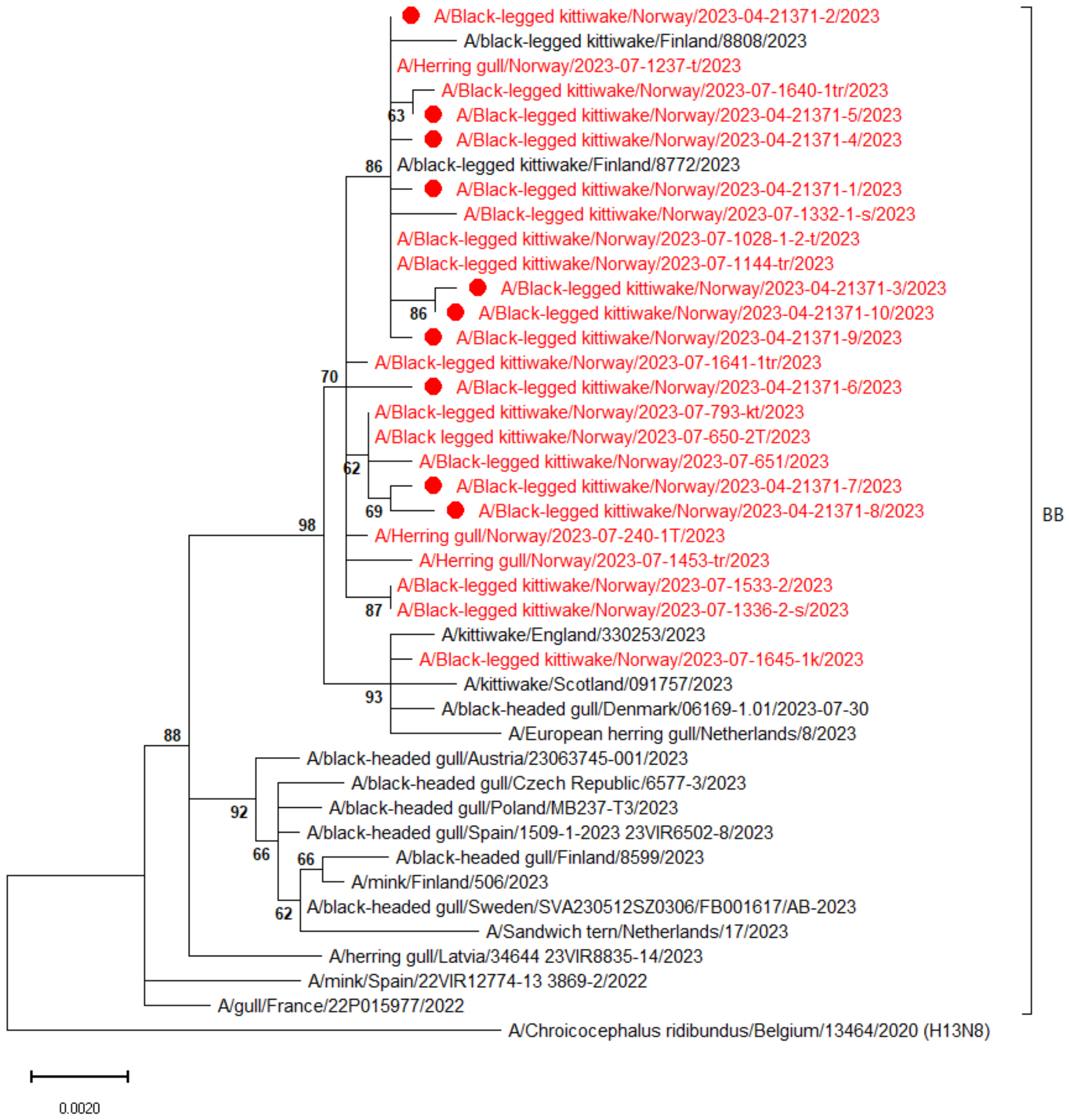
A/Black-legged kittiwake/Norway/2023-04-21371-2/2023
A/black-legged kittiwake/Finland/8808/2023
A/Herring gull/Norway/2023-07-1237-t/2023
A/Black-legged kittiwake/Norway/2023-07-1640-1tr/2023
A/Black-legged kittiwake/Norway/2023-04-21371-5/2023
A/Black-legged kittiwake/Norway/2023-04-21371-4/2023
A/black-legged kittiwake/Finland/8772/2023
A/Black-legged kittiwake/Norway/2023-04-21371-1/2023
A/Black-legged kittiwake/Norway/2023-07-1332-1-s/2023
A/Black-legged kittiwake/Norway/2023-07-1028-1-2-t/2023
A/Black-legged kittiwake/Norway/2023-07-1144-tr/2023
A/Black-legged kittiwake/Norway/2023-04-21371-3/2023
A/Black-legged kittiwake/Norway/2023-04-21371-10/2023
A/Black-legged kittiwake/Norway/2023-04-21371-9/2023
A/Black-legged kittiwake/Norway/2023-07-1641-1tr/2023
A/Black-legged kittiwake/Norway/2023-04-21371-6/2023
A/Black-legged kittiwake/Norway/2023-07-793-kt/2023
A/Black legged kittiwake/Norway/2023-07-650-2T/2023
A/Black-legged kittiwake/Norway/2023-07-651/2023
A/Black-legged kittiwake/Norway/2023-04-21371-7/2023
A/Black-legged kittiwake/Norway/2023-04-21371-8/2023
A/Herring gull/Norway/2023-07-240-1T/2023
A/Herring gull/Norway/2023-07-1453-tr/2023
A/Black-legged kittiwake/Norway/2023-07-1533-2/2023
A/Black-legged kittiwake/Norway/2023-07-1336-2-s/2023
A/kittiwake/England/330253/2023
A/Black-legged kittiwake/Norway/2023-07-1645-1k/2023
A/kittiwake/Scotland/091757/2023
A/black-headed gull/Denmark/06169-1.01/2023-07-30
A/European herring gull/Netherlands/8/2023
A/black-headed gull/Austria/23063745-001/2023
A/black-headed gull/Czech Republic/6577-3/2023
A/black-headed gull/Poland/MB237-T3/2023
A/black-headed gull/Spain/1509-1-2023 23VIR6502-8/2023
A/black-headed gull/Finland/8599/2023
A/mink/Finland/506/2023
A/black-headed gull/Sweden/SVA230512SZ0306/FB001617/AB-2023
A/Sandwich tern/Netherlands/17/2023
A/herring gull/Latvia/34644 23VIR8835-14/2023
A/mink/Spain/22VIR12774-13 3869-2/2022
A/gull/France/22P015977/2022
A/Chroicocephalus ridibundus/Belgium/13464/2020 (H13N8)
63
86
86
70
62
69
98
87
93
88
92
66
66
62
BB
0.0020

D)

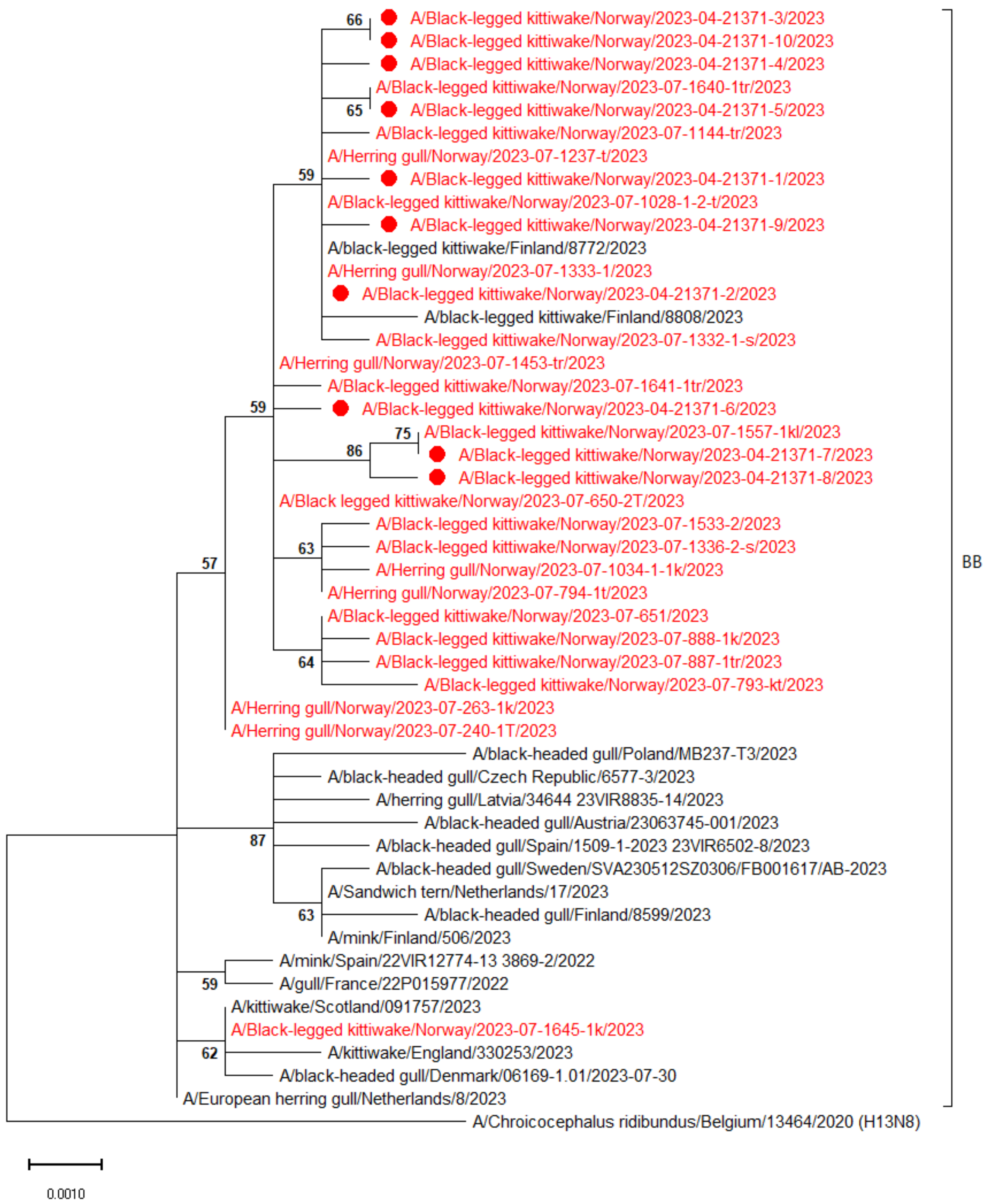
A/Black-legged kittiwake/Norway/2023-04-21371-3/2023
A/Black-legged kittiwake/Norway/2023-04-21371-10/2023
A/Black-legged kittiwake/Norway/2023-04-21371-4/2023
A/Black-legged kittiwake/Norway/2023-07-1640-1tr/2023
A/Black-legged kittiwake/Norway/2023-04-21371-5/2023
A/Black-legged kittiwake/Norway/2023-07-1144-tr/2023
A/Herring gull/Norway/2023-07-1237-t/2023
A/Black-legged kittiwake/Norway/2023-04-21371-1/2023
A/Black-legged kittiwake/Norway/2023-07-1028-1-2-t/2023
A/Black-legged kittiwake/Norway/2023-04-21371-9/2023
A/black-legged kittiwake/Finland/8772/2023
A/Herring gull/Norway/2023-07-1333-1/2023
A/Black-legged kittiwake/Norway/2023-04-21371-2/2023
A/black-legged kittiwake/Finland/8808/2023
A/Black-legged kittiwake/Norway/2023-07-1332-1-s/2023
A/Herring gull/Norway/2023-07-1453-tr/2023
A/Black-legged kittiwake/Norway/2023-07-1641-1tr/2023
A/Black-legged kittiwake/Norway/2023-04-21371-6/2023
A/Black-legged kittiwake/Norway/2023-07-1557-1kl/2023
A/Black-legged kittiwake/Norway/2023-04-21371-7/2023
A/Black-legged kittiwake/Norway/2023-04-21371-8/2023
A/Black legged kittiwake/Norway/2023-07-650-2T/2023
A/Black-legged kittiwake/Norway/2023-07-1533-2/2023
A/Black-legged kittiwake/Norway/2023-07-1336-2-s/2023
A/Herring gull/Norway/2023-07-1034-1-1k/2023
A/Herring gull/Norway/2023-07-794-1t/2023
A/Black-legged kittiwake/Norway/2023-07-651/2023
A/Black-legged kittiwake/Norway/2023-07-888-1k/2023
A/Black-legged kittiwake/Norway/2023-07-887-1tr/2023
A/Black-legged kittiwake/Norway/2023-07-793-kt/2023
A/Herring gull/Norway/2023-07-263-1k/2023
A/Herring gull/Norway/2023-07-240-1T/2023
A/black-headed gull/Poland/MB237-T3/2023
A/black-headed gull/Czech Republic/6577-3/2023
A/herring gull/Latvia/34644 23VIR8835-14/2023
A/black-headed gull/Austria/23063745-001/2023
A/black-headed gull/Spain/1509-1-2023 23VIR6502-8/2023
A/black-headed gull/Sweden/SVA230512SZ0306/FB001617/AB-2023
A/Sandwich tern/Netherlands/17/2023
A/black-headed gull/Finland/8599/2023
A/mink/Finland/506/2023
A/mink/Spain/22VIR12774-13 3869-2/2022
A/gull/France/22P015977/2022
A/kittiwake/Scotland/091757/2023
A/Black-legged kittiwake/Norway/2023-07-1645-1k/2023
A/kittiwake/England/330253/2023
A/black-headed gull/Denmark/06169-1.01/2023-07-30
A/European herring gull/Netherlands/8/2023
A/Chroicocephalus ridibundus/Belgium/13464/2020 (H13N8)
66
65
59
59
75
86
57
63
64
87
63
59
62
BB
0.0010

E)

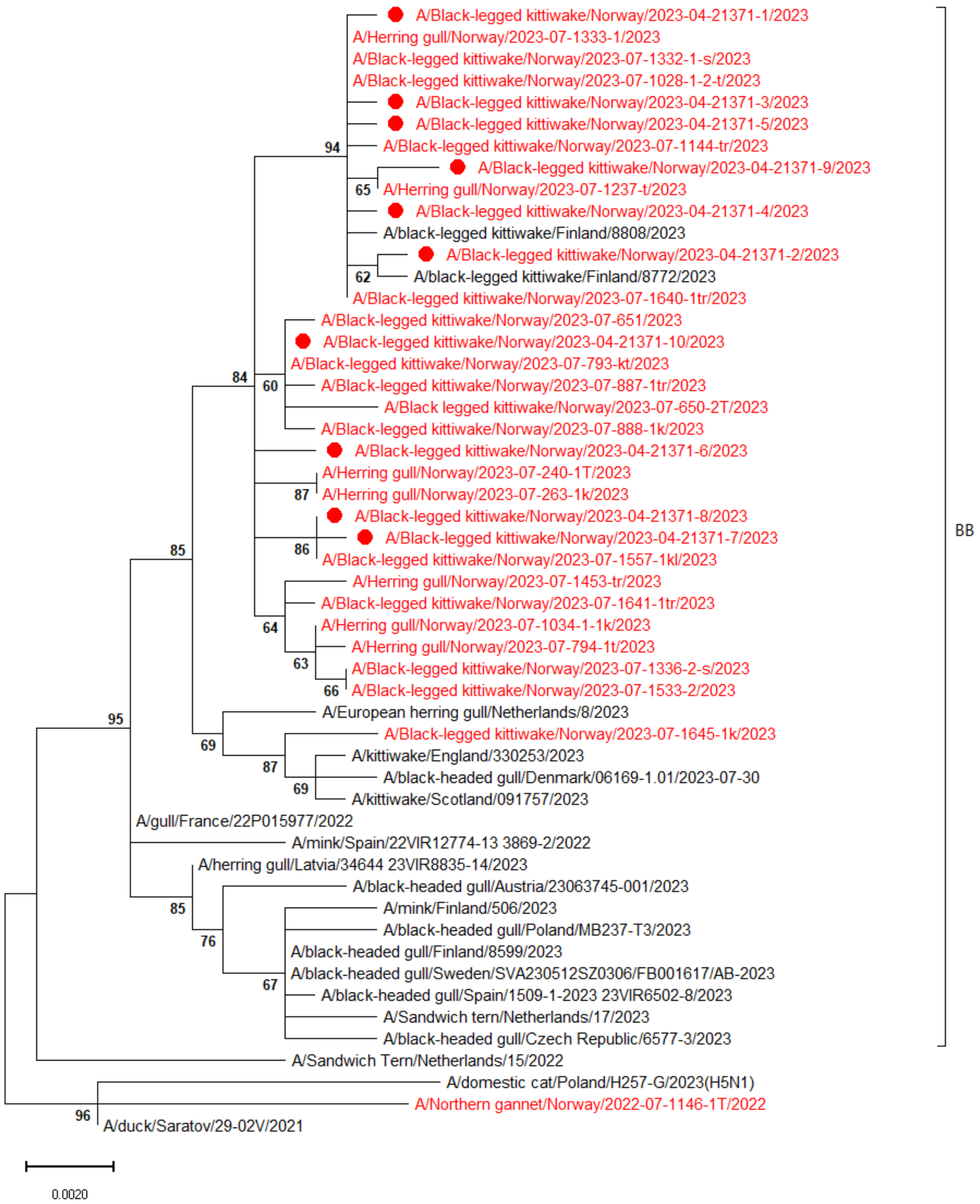
A/Black-legged kittiwake/Norway/2023-04-21371-1/2023
A/Herring gull/Norway/2023-07-1333-1/2023
A/Black-legged kittiwake/Norway/2023-07-1332-1-s/2023
A/Black-legged kittiwake/Norway/2023-07-1028-1-2-t/2023
A/Black-legged kittiwake/Norway/2023-04-21371-3/2023
A/Black-legged kittiwake/Norway/2023-04-21371-5/2023
94
A/Black-legged kittiwake/Norway/2023-07-1144-tr/2023
A/Black-legged kittiwake/Norway/2023-04-21371-9/2023
65
A/Herring gull/Norway/2023-07-1237-t/2023
A/Black-legged kittiwake/Norway/2023-04-21371-4/2023
A/black-legged kittiwake/Finland/8808/2023
A/Black-legged kittiwake/Norway/2023-04-21371-2/2023
62
A/black-legged kittiwake/Finland/8772/2023
A/Black-legged kittiwake/Norway/2023-07-1640-1tr/2023
A/Black-legged kittiwake/Norway/2023-07-651/2023
A/Black-legged kittiwake/Norway/2023-04-21371-10/2023
A/Black-legged kittiwake/Norway/2023-07-793-kt/2023
84
60
A/Black-legged kittiwake/Norway/2023-07-887-1tr/2023
A/Black legged kittiwake/Norway/2023-07-650-2T/2023
A/Black-legged kittiwake/Norway/2023-07-888-1k/2023
A/Black-legged kittiwake/Norway/2023-04-21371-6/2023
A/Herring gull/Norway/2023-07-240-1T/2023
87
A/Herring gull/Norway/2023-07-263-1k/2023
A/Black-legged kittiwake/Norway/2023-04-21371-8/2023
BB
A/Black-legged kittiwake/Norway/2023-04-21371-7/2023
85
86
A/Black-legged kittiwake/Norway/2023-07-1557-1kl/2023
A/Herring gull/Norway/2023-07-1453-tr/2023
A/Black-legged kittiwake/Norway/2023-07-1641-1tr/2023
64
A/Herring gull/Norway/2023-07-1034-1-1k/2023
A/Herring gull/Norway/2023-07-794-1t/2023
63
A/Black-legged kittiwake/Norway/2023-07-1336-2-s/2023
66
A/Black-legged kittiwake/Norway/2023-07-1533-2/2023
A/European herring gull/Netherlands/8/2023
95
A/Black-legged kittiwake/Norway/2023-07-1645-1k/2023
69
A/kittiwake/England/330253/2023
87
A/black-headed gull/Denmark/06169-1.01/2023-07-30
69
A/kittiwake/Scotland/091757/2023
A/gull/France/22P015977/2022
A/mink/Spain/22VIR12774-13 3869-2/2022
A/herring gull/Latvia/34644 23VIR8835-14/2023
A/black-headed gull/Austria/23063745-001/2023
85
A/mink/Finland/506/2023
76
A/black-headed gull/Poland/MB237-T3/2023
A/black-headed gull/Finland/8599/2023
A/black-headed gull/Sweden/SVA230512SZ0306/FB001617/AB-2023
67
A/black-headed gull/Spain/1509-1-2023 23VIR6502-8/2023
A/Sandwich tern/Netherlands/17/2023
A/black-headed gull/Czech Republic/6577-3/2023
A/Sandwich Tern/Netherlands/15/2022
A/domestic cat/Poland/H257-G/2023(H5N1)
A/Northern gannet/Norway/2022-07-1146-1T/2022
96
A/duck/Saratov/29-02V/2021
0.0020

F)

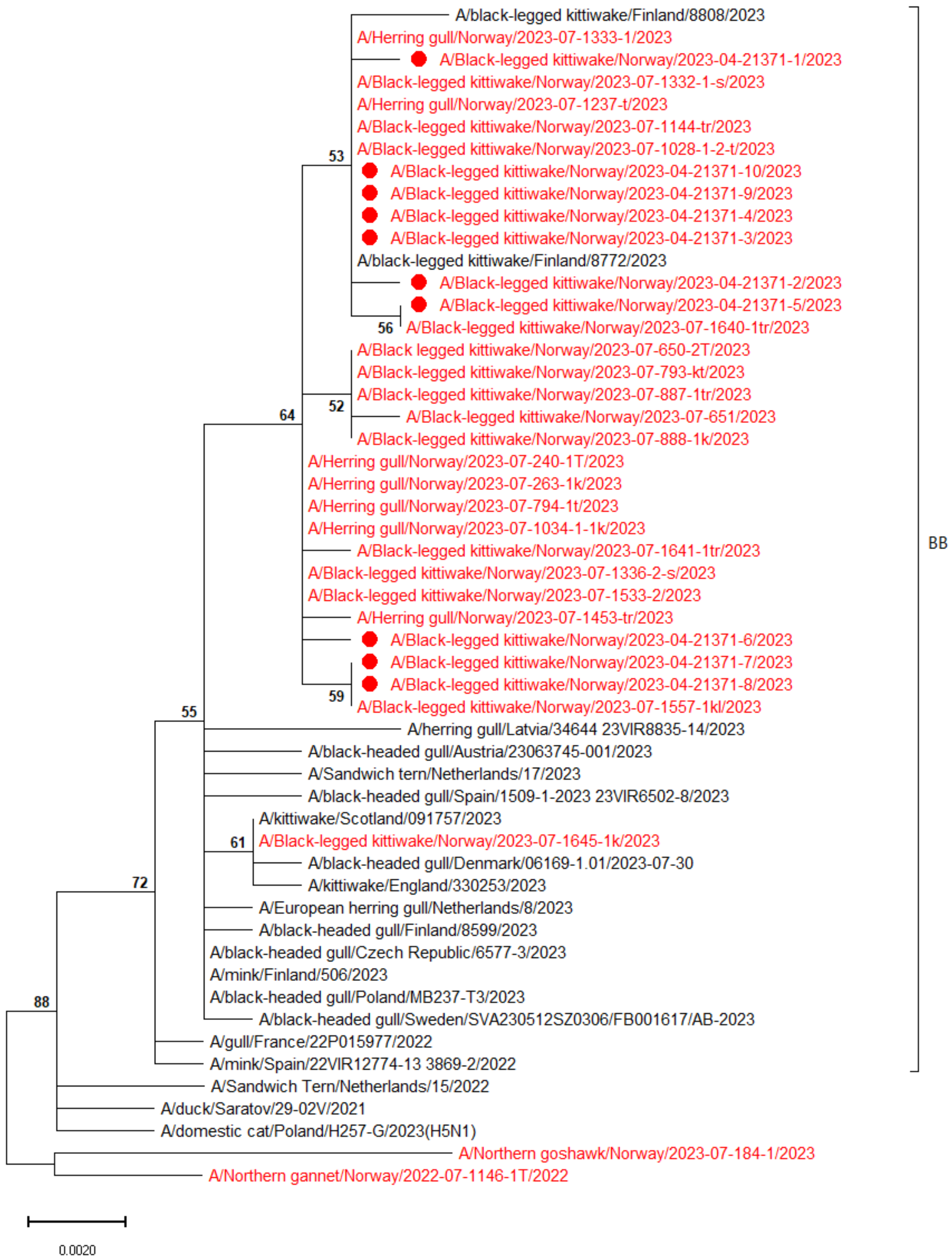
A/black-legged kittiwake/Finland/8808/2023
A/Herring gull/Norway/2023-07-1333-1/2023
A/Black-legged kittiwake/Norway/2023-04-21371-1/2023
A/Black-legged kittiwake/Norway/2023-07-1332-1-s/2023
A/Herring gull/Norway/2023-07-1237-t/2023
A/Black-legged kittiwake/Norway/2023-07-1144-tr/2023
53
A/Black-legged kittiwake/Norway/2023-07-1028-1-2-t/2023
A/Black-legged kittiwake/Norway/2023-04-21371-10/2023
A/Black-legged kittiwake/Norway/2023-04-21371-9/2023
A/Black-legged kittiwake/Norway/2023-04-21371-4/2023
A/Black-legged kittiwake/Norway/2023-04-21371-3/2023
A/black-legged kittiwake/Finland/8772/2023
A/Black-legged kittiwake/Norway/2023-04-21371-2/2023
A/Black-legged kittiwake/Norway/2023-04-21371-5/2023
56
A/Black-legged kittiwake/Norway/2023-07-1640-1tr/2023
A/Black legged kittiwake/Norway/2023-07-650-2T/2023
A/Black-legged kittiwake/Norway/2023-07-793-kt/2023
A/Black-legged kittiwake/Norway/2023-07-887-1tr/2023
52
64
A/Black-legged kittiwake/Norway/2023-07-651/2023
A/Black-legged kittiwake/Norway/2023-07-888-1k/2023
A/Herring gull/Norway/2023-07-240-1T/2023
A/Herring gull/Norway/2023-07-263-1k/2023
A/Herring gull/Norway/2023-07-794-1t/2023
A/Herring gull/Norway/2023-07-1034-1-1k/2023
A/Black-legged kittiwake/Norway/2023-07-1641-1tr/2023
A/Black-legged kittiwake/Norway/2023-07-1336-2-s/2023
A/Black-legged kittiwake/Norway/2023-07-1533-2/2023
A/Herring gull/Norway/2023-07-1453-tr/2023
A/Black-legged kittiwake/Norway/2023-04-21371-6/2023
A/Black-legged kittiwake/Norway/2023-04-21371-7/2023
A/Black-legged kittiwake/Norway/2023-04-21371-8/2023
59
A/Black-legged kittiwake/Norway/2023-07-1557-1kl/2023
55
A/herring gull/Latvia/34644 23VIR8835-14/2023
A/black-headed gull/Austria/23063745-001/2023
A/Sandwich tern/Netherlands/17/2023
A/black-headed gull/Spain/1509-1-2023 23VIR6502-8/2023
A/kittiwake/Scotland/091757/2023
61
A/Black-legged kittiwake/Norway/2023-07-1645-1k/2023
A/black-headed gull/Denmark/06169-1.01/2023-07-30
A/kittiwake/England/330253/2023
72
A/European herring gull/Netherlands/8/2023
A/black-headed gull/Finland/8599/2023
A/black-headed gull/Czech Republic/6577-3/2023
A/mink/Finland/506/2023
88
A/black-headed gull/Poland/MB237-T3/2023
A/black-headed gull/Sweden/SVA230512SZ0306/FB001617/AB-2023
A/gull/France/22P015977/2022
A/mink/Spain/22VIR12774-13 3869-2/2022
A/Sandwich Tern/Netherlands/15/2022
A/duck/Saratov/29-02V/2021
A/domestic cat/Poland/H257-G/2023(H5N1)
A/Northern goshawk/Norway/2023-07-184-1/2023
A/Northern gannet/Norway/2022-07-1146-1T/2022
BB
0.0020

G)

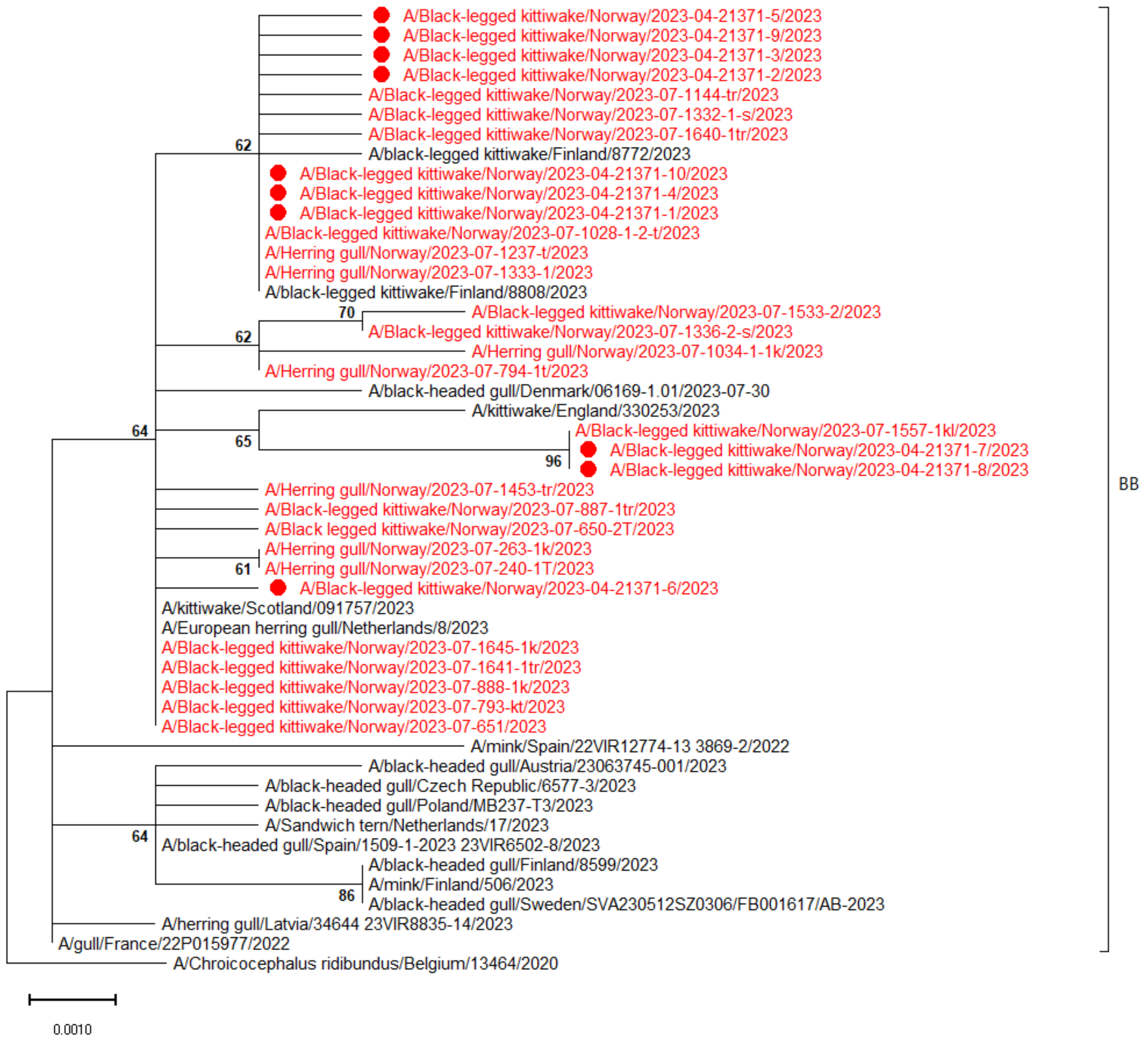


Figure 2. Midpoint rooted phylogenetic trees showing the genetic relationship between ten HPAI H5N1 viruses identified in dead Black-legged Kittiwakes from Ekkerøy, Norway, in 2023 (red dots), and contemporary viruses from Norway (red) and Europe (black), including the coding part of the gene segments A) PB2, B) PB1, C) PA, D) NP, E) NA, F) M, G) NS. Bootstrap values >50 are shown.
