## Appendix 2 for "Highly Pathogenic Avian Influenza A(H5N1) Caused Mass Death among Black-legged Kittiwakes (*Rissa tridactyla*) in Norway, 2023"

**Appendix 2 Table 1. Data included in the phylogenetic analyses.**

We thank the authors, originating and submitting laboratories of the sequences from GISAID's EpiFlu Database (www.gisaid.org) on which this research is based. A detailed list is found below.

| GISAID isolat acc.no | Isolate name (ID, label) | Pathogenicy | Subtype | Clade | Genotype | PB2 | PB1 | PA | HA | NP | NA | M | NS | Collection date |
|---|---|---|---|---|---|---|---|---|---|---|---|---|---|---|
| EPI ISL 18292959 | A/black-legged kittiwake/Finland/8808/2023 | HPAI | H5N1 | 2.3.4.4b | BB | EPI2755467 | EPI2755468 | EPI2755466 | EPI2755470 | EPI2755463 | EPI2755469 | EPI2755465 | EPI2755464 | 2023-07-28 |
| EPI ISL 18728075 | A/black-legged kittiwake/Finland/8772/2023 | HPAI | H5N1 | 2.3.4.4b | BB | EPI2892377 | EPI2892379 | EPI2892381 | EPI2892391 | EPI2892421 | EPI2892446 | EPI2892471 | EPI2892493 | 2023-07-21 |
| EPI ISL 18755127 | A/black-headed gull/Finland/8599/2023 | HPAI | H5N1 | 2.3.4.4b | BB | EPI2907863 | EPI2907864 | EPI2907865 | EPI2907866 | EPI2907867 | EPI2907868 | EPI2907869 | EPI2907870 | 2023-07-20 |
| EPI ISL 18271510 | A/black-headed gull/Denmark/06169-1.01/2023-07-3 | HPAI | H5N1 | 2.3.4.4b | BB | EPI2745873 | EPI2745874 | EPI2745875 | EPI2745876 | EPI2745877 | EPI2745878 | EPI2745879 | EPI2745880 | 2023-07-30 |
| EPI ISL 13778522 | A/Sandwich Tern/Netherlands/15/2022 | HPAI | H5N2 | 2.3.4.4b | AB | EPI2088979 | EPI2088980 | Excluded* | EPI2088982 | Excluded* | EPI2088981 | EPI2088977 | Excluded* | 2022-06-24 |
| EPI ISL 18882069 | A/Sandwich tern/Netherlands/17/2023 | HPAI | H5N3 | 2.3.4.4b | BB | EPI3010420 | EPI3010421 | EPI3010419 | EPI3010423 | EPI3010416 | EPI3010422 | EPI3010418 | EPI3010417 | 2023-07-04 |
| EPI ISL 17267156 | A/European herring gull/Netherlands/8/2023 | HPAI | H5N1 | 2.3.4.4b | BB | EPI2475220 | EPI2475221 | EPI2475219 | EPI2475223 | EPI2475216 | EPI2475222 | EPI2475218 | EPI2475217 | 2023-03-03 |
| EPI ISL 18112333 | A/kittiwake/England/330253/2023 | HPAI | H5N1 | 2.3.4.4b | BB | EPI2691234 | EPI2691235 | EPI2691233 | EPI2691237 | EPI2691230 | EPI2691236 | EPI2691232 | EPI2691231 | 2023-07-08 |
| EPI ISL 18112336 | A/kittiwake/Scotland/091757/2023 | HPAI | H5N1 | 2.3.4.4b | BB | EPI2691258 | EPI2691259 | EPI2691257 | EPI2691261 | EPI2691254 | EPI2691260 | EPI2691256 | EPI2691255 | 2023-06-30 |
| EPI ISL 18075732 | A/black-headed gull/Spain/1509-1-2023 23VIR6502-8/2023 | HPAI | H5N1 | 2.3.4.4b | BB | EPI2671555 | EPI2671556 | EPI2671554 | EPI2671558 | EPI2671551 | EPI2671557 | EPI2671553 | EPI2671552 | 2023-03-16 |
| EPI ISL 18058689 | A/black-headed gull/Austria/23063745-001/2023 | HPAI | H5N1 | 2.3.4.4b | BB | EPI2664923 | EPI2664924 | EPI2664922 | EPI2664926 | EPI2664919 | EPI2664925 | EPI2664921 | EPI2664920 | 2023-05-12 |
| EPI ISL 18033241 | A/black-headed gull/Poland/MB237-T3/2023 | HPAI | H5N1 | 2.3.4.4b | BB | EPI2652213 | EPI2652214 | EPI2652212 | EPI2648247 | EPI2652209 | EPI2652215 | EPI2652211 | EPI2652210 | 2023-05-30 |
| EPI ISL 17716812 | A/black-headed gull/Czech Republic/6577-3/2023 | HPAI | H5N1 | 2.3.4.4b | BB | EPI2577985 | EPI2577986 | EPI2577984 | EPI2577988 | EPI2577981 | EPI2577987 | EPI2577983 | EPI2577982 | 2023-04-24 |
| EPI ISL 7622539 | A/Chroicocephalus ridibundus/Belgium/13464/2020 | LPAI | H13N8 | 2.3.4.4b | BB | Excluded* | Excluded* | EPI1942889 | Excluded* | EPI1942891 | Excluded* | Excluded* | EPI1942894 | 2021-12-09 |
| EPI ISL 13519451 | A/gull/France/22P015977/2022 | HPAI | H5N1 | 2.3.4.4b | BB | EPI2078351 | EPI2078352 | EPI2078353 | EPI2078354 | EPI2078355 | EPI2078356 | EPI2078357 | EPI2078358 | 2022-05-11 |
| EPI ISL 5463797 | A/duck/Saratov/29-02V/2021 | HPAI | H5N1 | 2.3.4.4b | AB | EPI1922922 | EPI1922923 | Excluded* | EPI1922925 | Excluded* | EPI1922924 | EPI1922920 | Excluded* | 2021-09-30 |
| EPI ISL 18455202 | A/Northern gannet/Norway/2022-07-1146-1T/2022 | HPAI | H5N1 | 2.3.4.4b | C | Excluded* | EPI2782075 | Excluded* | EPI2782077 | Excluded* | EPI2782079 | EPI2782080 | Excluded* | 2022-06-16 |
| EPI ISL 18851332 | A/Northern goshawk/Norway/2023-07-184-1/2023 | HPAI | H5N5 | 2.3.4.4b | I | Excluded* | Excluded* | Excluded* | EPI2972840 | Excluded* | Excluded** | EPI2972843 | Excluded* | 2023-03-13 |
| EPI ISL 17971991 | A/domestic cat/Poland/H257-G/2023(H5N1) | HPAI | H5N1 | 2.3.4.4b | CH | EPI2616171 | EPI2616172 | Excluded* | EPI2616174 | Excluded* | EPI2616173 | EPI2616169 | Excluded* | 2023-06-24 |
| EPI ISL 18292952 | A/mink/Finland/506/2023 | HPAI | H5N1 | 2.3.4.4b | BB | EPI2755418 | EPI2755417 | EPI2755416 | EPI2755411 | EPI2755414 | EPI2755413 | EPI2755412 | EPI2755415 | 2023-07-03 |
| EPI ISL 16507095 | A/mink/Spain/22VIR12774-13 3869-2/2022 | HPAI | H5N1 | 2.3.4.4b | BB | EPI2291507 | EPI2291508 | EPI2291506 | EPI2291510 | EPI2291503 | EPI2291509 | EPI2291505 | EPI2291504 | 2022-10-26 |
| EPI ISL 17787071 | A/black-headed gull/Sweden/SVA230512SZ0306/FB001617/AB-2 | HPAI | H5N1 | 2.3.4.4b | BB | EPI2589386 | EPI2589387 | EPI2589388 | EPI2589389 | EPI2589390 | EPI2589391 | EPI2589392 | EPI2589393 | 2023-05-11 |
| EPI ISL 18509067 | A/herring gull/Latvia/34644 23VIR8835-14/2023 | HPAI | H5N1 | 2.3.4.4b | BB | EPI2794089 | EPI2794090 | EPI2794088 | EPI2794092 | EPI2794085 | EPI2794091 | EPI2794087 | EPI2794086 | 2023-05-29 |
| EPI ISL 18455264 | A/Black legged kittiwake/Norway/2023-07-650-2T/2023 | HPAI | H5N1 | 2.3.4.4b | BB | EPI2782185 | EPI2782186 | EPI2782187 | EPI2782188 | EPI2782189 | EPI2782190 | EPI2782191 | EPI2782192 | 2023-05-23 |
| EPI ISL 18939118 | A/Herring gull/Norway/2023-07-240-1T/2023 | HPAI | H5N1 | 2.3.4.4b | BB | EPI3077700 | EPI3077701 | EPI3077699 | EPI3077703 | EPI3077696 | EPI3077702 | EPI3077698 | EPI3077697 | 2023-04-12 |
| EPI ISL 18939125 | A/Herring gull/Norway/2023-07-263-1k/2023 | HPAI | H5N1 | 2.3.4.4b | BB | missing | missing | missing | EPI3077748 | EPI3077744 | EPI3077746 | EPI3077746 | EPI3077745 | 2023-04-18 |
| EPI ISL 18939119 | A/Black-legged kittiwake/Norway/2023-07-651/2023 | HPAI | H5N1 | 2.3.4.4b | BB | EPI3077708 | EPI3077709 | EPI3077707 | EPI3077711 | EPI3077704 | EPI3077710 | EPI3077706 | EPI3077705 | 2023-05-16 |
| EPI ISL 18939120 | A/Black-legged kittiwake/Norway/2023-07-793-kt/2023 | HPAI | H5N1 | 2.3.4.4b | BB | EPI3077716 | EPI3077717 | EPI3077715 | EPI3077719 | EPI3077712 | EPI3077718 | EPI3077714 | EPI3077713 | 2023-05-25 |
| EPI ISL 18939126 | A/Herring gull/Norway/2023-07-794-1t/2023 | HPAI | H5N1 | 2.3.4.4b | BB | EPI3077752 | Excluded* | missing | EPI3077755 | EPI3077749 | EPI3077754 | EPI3077751 | EPI3077750 | 2023-05-26 |
| EPI ISL 18939121 | A/Black-legged kittiwake/Norway/2023-07-887-1tr/2023 | HPAI | H5N1 | 2.3.4.4b | BB | EPI3077724 | EPI3077725 | Excluded* | EPI3077727 | EPI3077720 | EPI3077726 | EPI3077722 | EPI3077721 | 2023-05-25 |
| EPI ISL 18939122 | A/Black-legged kittiwake/Norway/2023-07-888-1k/2023 | HPAI | H5N1 | 2.3.4.4b | BB | EPI3077732 | EPI3077733 | Excluded* | EPI3077735 | EPI3077728 | EPI3077734 | EPI3077730 | EPI3077729 | 2023-06-02 |
| EPI ISL 18939106 | A/Black-legged kittiwake/Norway/2023-07-1028-1-2-t/2023 | HPAI | H5N1 | 2.3.4.4b | BB | EPI3077604 | EPI3077605 | EPI3077603 | EPI3077607 | EPI3077600 | EPI3077606 | EPI3077602 | EPI3077601 | 2023-06-07 |
| EPI ISL 18939107 | A/Herring gull/Norway/2023-07-1034-1-1k/2023 | HPAI | H5N1 | 2.3.4.4b | BB | EPI3077612 | Excluded* | Excluded* | EPI3077615 | EPI3077608 | EPI3077614 | EPI3077610 | EPI3077609 | 2023-06-15 |
| EPI ISL 18939108 | A/Black-legged kittiwake/Norway/2023-07-1144-tr/2023 | HPAI | H5N1 | 2.3.4.4b | BB | EPI3077620 | EPI3077621 | EPI3077619 | EPI3077623 | EPI3077616 | EPI3077622 | EPI3077618 | EPI3077617 | 2023-06-15 |
| EPI ISL 18939109 | A/Herring gull/Norway/2023-07-1237-t/2023 | HPAI | H5N1 | 2.3.4.4b | BB | EPI3077628 | EPI3077629 | EPI3077627 | EPI3077631 | EPI3077624 | EPI3077630 | EPI3077626 | EPI3077625 | 2023-06-22 |
| EPI ISL 18939110 | A/Black-legged kittiwake/Norway/2023-07-1332-1-s/2023 | HPAI | H5N1 | 2.3.4.4b | BB | EPI3077636 | EPI3077637 | EPI3077635 | EPI3077639 | EPI3077632 | EPI3077638 | EPI3077634 | EPI3077633 | 2023-07-13 |
| EPI ISL 18939124 | A/Herring gull/Norway/2023-07-1333-1/2023 | HPAI | H5N1 | 2.3.4.4b | BB | EPI3077740 | Excluded* | missing | EPI3077743 | EPI3077737 | EPI3077742 | EPI3077739 | EPI3077738 | 2023-06-28 |
| EPI ISL 18939111 | A/Black-legged kittiwake/Norway/2023-07-1336-2-s/2023 | HPAI | H5N1 | 2.3.4.4b | BB | EPI3077644 | EPI3077645 | EPI3077643 | EPI3077647 | EPI3077640 | EPI3077646 | EPI3077642 | EPI3077641 | 2023-07-04 |
| EPI ISL 18939112 | A/Herring gull/Norway/2023-07-1453-tr/2023 | HPAI | H5N1 | 2.3.4.4b | BB | Excluded* | EPI3077653 | EPI3077651 | EPI3077655 | EPI3077648 | EPI3077654 | EPI3077650 | EPI3077649 | 2023-07-05 |
| EPI ISL 18939113 | A/Black-legged kittiwake/Norway/2023-07-1533-2/2023 | HPAI | H5N1 | 2.3.4.4b | BB | EPI3077660 | EPI3077661 | EPI3077659 | EPI3077663 | EPI3077656 | EPI3077662 | EPI3077658 | EPI3077657 | 2023-07-19 |
| EPI ISL 18939114 | A/Black-legged kittiwake/Norway/2023-07-1557-1kl/2023 | HPAI | H5N1 | 2.3.4.4b | BB | EPI3077668 | EPI3077669 | Excluded* | EPI3077671 | EPI3077664 | EPI3077670 | EPI3077666 | EPI3077665 | 2023-07-20 |
| EPI ISL 18939115 | A/Black-legged kittiwake/Norway/2023-07-1640-1tr/2023 | HPAI | H5N1 | 2.3.4.4b | BB | EPI3077676 | EPI3077677 | EPI3077675 | EPI3077679 | EPI3077672 | EPI3077678 | EPI3077674 | EPI3077673 | 2023-07-20 |
| EPI ISL 18939116 | A/Black-legged kittiwake/Norway/2023-07-1641-1tr/2023 | HPAI | H5N1 | 2.3.4.4b | BB | EPI3077684 | EPI3077685 | EPI3077683 | EPI3077687 | EPI3077680 | EPI3077686 | EPI3077682 | EPI3077681 | 2023-07-20 |
| EPI ISL 18939117 | A/Black-legged kittiwake/Norway/2023-07-1645-1k/2023 | HPAI | H5N1 | 2.3.4.4b | BB | EPI3077692 | EPI3077693 | EPI3077691 | EPI3077695 | EPI3077688 | EPI3077694 | EPI3077690 | EPI3077689 | 2023-07-20 |
| EPI ISL 18939097 | A/Black-legged kittiwake/Norway/2023-04-21371-1/2023 | HPAI | H5N1 | 2.3.4.4b | BB | EPI3077532 | EPI3077533 | EPI3077531 | EPI3077535 | EPI3077528 | EPI3077534 | EPI3077530 | EPI3077529 | 2023-07-26 |
| EPI ISL 18939098 | A/Black-legged kittiwake/Norway/2023-04-21371-2/2023 | HPAI | H5N1 | 2.3.4.4b | BB | EPI3077540 | EPI3077541 | EPI3077539 | EPI3077543 | EPI3077536 | EPI3077542 | EPI3077538 | EPI3077537 | 2023-07-26 |
| EPI ISL 18939099 | A/Black-legged kittiwake/Norway/2023-04-21371-3/2023 | HPAI | H5N1 | 2.3.4.4b | BB | EPI3077548 | EPI3077549 | EPI3077547 | EPI3077551 | EPI3077544 | EPI3077550 | EPI3077546 | EPI3077545 | 2023-07-26 |
| EPI ISL 18939100 | A/Black-legged kittiwake/Norway/2023-04-21371-4/2023 | HPAI | H5N1 | 2.3.4.4b | BB | EPI3077556 | EPI3077557 | EPI3077555 | EPI3077559 | EPI3077552 | EPI3077558 | EPI3077554 | EPI3077553 | 2023-07-26 |
| EPI ISL 18939101 | A/Black-legged kittiwake/Norway/2023-04-21371-5/2023 | HPAI | H5N1 | 2.3.4.4b | BB | EPI3077564 | EPI3077565 | EPI3077563 | EPI3077567 | EPI3077560 | EPI3077566 | EPI3077562 | EPI3077561 | 2023-07-26 |
| EPI ISL 18939102 | A/Black-legged kittiwake/Norway/2023-04-21371-6/2023 | HPAI | H5N1 | 2.3.4.4b | BB | EPI3077572 | EPI3077573 | EPI3077571 | EPI3077575 | EPI3077568 | EPI3077574 | EPI3077570 | EPI3077569 | 2023-07-26 |
| EPI ISL 18939103 | A/Black-legged kittiwake/Norway/2023-04-21371-7/2023 | HPAI | H5N1 | 2.3.4.4b | BB | EPI3077580 | EPI3077581 | EPI3077579 | EPI3077583 | EPI3077576 | EPI3077582 | EPI3077578 | EPI3077577 | 2023-07-26 |
| EPI ISL 18939104 | A/Black-legged kittiwake/Norway/2023-04-21371-8/2023 | HPAI | H5N1 | 2.3.4.4b | BB | EPI3077588 | EPI3077589 | EPI3077587 | EPI3077591 | EPI3077584 | EPI3077590 | EPI3077586 | EPI3077585 | 2023-07-26 |
| EPI ISL 18939105 | A/Black-legged kittiwake/Norway/2023-04-21371-9/2023 | HPAI | H5N1 | 2.3.4.4b | BB | EPI3077596 | EPI3077597 | EPI3077595 | EPI3077599 | EPI3077592 | EPI3077598 | EPI3077594 | EPI3077593 | 2023-07-26 |
| EPI ISL 18939096 | A/Black-legged kittiwake/Norway/2023-04-21371-10/2023 | HPAI | H5N1 | 2.3.4.4b | BB | EPI3077524 | EPI3077525 | EPI3077523 | EPI3077527 | EPI3077520 | EPI3077526 | EPI3077522 | EPI3077521 | 2023-07-26 |

*Sequence excluded due to low or too large distance
**N5 sequence excluded

| Collection montl | Collection year | Continent/region | Country | County (NO only) | Muncicipality (NO only) | Species (English) | Species (latin) | Family | Order | Originating lab |
|---|---|---|---|---|---|---|---|---|---|---|
| July | 2023 | Nordic | Finland | | | Black-legged kittiwake | *Rissa tridactyla* | *Laridae* | Charadriiformes | Finnish Food Authority |
| July | 2023 | Nordic | Finland | | | Black-legged kittiwake | *Rissa tridactyla* | *Laridae* | Charadriiformes | Finnish Food Authority |
| July | 2023 | Nordic | Finland | | | Black-headed gull | *Chroicocephalus ridibundus* | *Laridae* | Charadriiformes | Finnish Food Authority |
| July | 2023 | Nordic | Denmark | | | Black-headed gull | *Chroicocephalus ridibundus* | *Laridae* | Charadriiformes | Statens Serum Institute |
| June | 2022 | Other European | Netherlands | | | Sandwich tern | *Thalasseus sandvicensis* | *Laridae* | Charadriiformes | Erasmus Medical Center |
| July | 2023 | Other European | Netherlands | | | Sandwich tern | *Thalasseus sandvicensis* | *Laridae* | Charadriiformes | Erasmus Medical Center |
| March | 2023 | Other European | Netherlands | | | Herring gull | *Larus argentatus* | *Laridae* | Charadriiformes | Erasmus Medical Center |
| July | 2023 | Other European | England | | | Black-legged kittiwake | *Rissa tridactyla* | *Laridae* | Charadriiformes | Animal and Plant Health Agency (APHA) |
| June | 2023 | Other European | Scotland | | | Black-legged kittiwake | *Rissa tridactyla* | *Laridae* | Charadriiformes | Animal and Plant Health Agency (APHA) |
| March | 2023 | Other European | Spania | | | Black-headed gull | *Chroicocephalus ridibundus* | *Laridae* | Charadriiformes | Laboratorio Central de Veterinaria |
| May | 2023 | Other European | Austria | | | Black-headed gull | *Chroicocephalus ridibundus* | *Laridae* | Charadriiformes | Austrian Agency for Health and Food Safety ( |
| May | 2023 | Other European | Poland | | | Black-headed gull | *Chroicocephalus ridibundus* | *Laridae* | Charadriiformes | National Veterinary Research Institut Poland, I |
| April | 2023 | Other European | Czech Republic | | | Black-headed gull | *Chroicocephalus ridibundus* | *Laridae* | Charadriiformes | State Veterinary Institute Prague |
| December | 2021 | Other European | Belgium | | | Black-headed gull | *Chroicocephalus ridibundus* | *Laridae* | Charadriiformes | Sciensano - Animal Infectious Diseases |
| May | 2022 | Other European | France | | | Gull | *Larus sp.* | *Laridae* | Charadriiformes | Anses (Ploufragan-Plouzané) |
| September | 2021 | Other European | Saratov | | | Domestic duck | *Anas platyrhynchos f. domestica* | *Anatidae* | Anseriformes | Center of Hygiene and Epidemiology in Sarato |
| June | 2022 | Nordic | Norway | Hordaland | Bergen | Northern gannet | *Morus bassanus* | *Sulidae* | Suliformes | Norwegian Veterinary Institute |
| March | 2023 | Nordic | Norway | Rogaland | Stavanger | Northern goshawk | *Accipiter gentilis* | *Accipitridae* | Accipitriformes | Norwegian Veterinary Institute |
| June | 2023 | Other European | Poland | | | Domestic cat | *Felis catus* | *Felidae* | Carnivora | National Veterinary Research Institut Poland, I |
| July | 2023 | Nordic | Finland | | | American mink | *Neovison vison* | *Mustelidae* | Carnivora | Finnish Food Authority |
| October | 2022 | Other European | Spain | | | American mink | *Neovison vison* | *Mustelidae* | Carnivora | Istituto Zooprofilattico Sperimentale delle Ven |
| May | 2023 | Nordic | Sweden | | | Black-headed gull | *Chroicocephalus ridibundus* | *Laridae* | Charadriiformes | Swedish Veterinary Agency (SVA) |
| May | 2023 | Other European | Lativa | | | Herring gull | *Larus argentatus* | *Laridae* | Charadriiformes | Institute of Food Safety, Animal Health and En |
| May | 2023 | Nordic | Norway | Troms | Harstad | Black-legged kittiwake | *Rissa tridactyla* | *Laridae* | Charadriiformes | Norwegian Veterinary Institute |
| April | 2023 | Nordic | Norway | Vestland | Bergen | Herring gull | *Larus argentatus* | *Laridae* | Charadriiformes | Norwegian Veterinary Institute |
| April | 2023 | Nordic | Norway | Vestland | Bjørnafjorden | Herring gull | *Larus argentatus* | *Laridae* | Charadriiformes | Norwegian Veterinary Institute |
| May | 2023 | Nordic | Norway | Troms | Salangen | Black-legged kittiwake | *Rissa tridactyla* | *Laridae* | Charadriiformes | Norwegian Veterinary Institute |
| May | 2023 | Nordic | Norway | Troms | Senja | Black-legged kittiwake | *Rissa tridactyla* | *Laridae* | Charadriiformes | Norwegian Veterinary Institute |
| May | 2023 | Nordic | Norway | Nordland | Moskenes | Herring gull | *Larus argentatus* | *Laridae* | Charadriiformes | Norwegian Veterinary Institute |
| May | 2023 | Nordic | Norway | Troms | Kvæfjord | Black-legged kittiwake | *Rissa tridactyla* | *Laridae* | Charadriiformes | Norwegian Veterinary Institute |
| June | 2023 | Nordic | Norway | Finnmark | Hammerfest | Black-legged kittiwake | *Rissa tridactyla* | *Laridae* | Charadriiformes | Norwegian Veterinary Institute |
| June | 2023 | Nordic | Norway | Finnmark | Berlevåg | Black-legged kittiwake | *Rissa tridactyla* | *Laridae* | Charadriiformes | Norwegian Veterinary Institute |
| June | 2023 | Nordic | Norway | Troms | Tromsø | Herring gull | *Larus argentatus* | *Laridae* | Charadriiformes | Norwegian Veterinary Institute |
| June | 2023 | Nordic | Norway | Finnmark | Båtsfjord | Black-legged kittiwake | *Rissa tridactyla* | *Laridae* | Charadriiformes | Norwegian Veterinary Institute |
| June | 2023 | Nordic | Norway | Finnmark | Vardø | Herring gull | *Larus argentatus* | *Laridae* | Charadriiformes | Norwegian Veterinary Institute |
| July | 2023 | Nordic | Norway | Finnmark | Båtsfjord | Black-legged kittiwake | *Rissa tridactyla* | *Laridae* | Charadriiformes | Norwegian Veterinary Institute |
| June | 2023 | Nordic | Norway | Rogaland | Karmøy | Herring gull | *Larus argentatus* | *Laridae* | Charadriiformes | Norwegian Veterinary Institute |
| July | 2023 | Nordic | Norway | Finnmark | Alta | Black-legged kittiwake | *Rissa tridactyla* | *Laridae* | Charadriiformes | Norwegian Veterinary Institute |
| July | 2023 | Nordic | Norway | Nordland | Bodø | Herring gull | *Larus argentatus* | *Laridae* | Charadriiformes | Norwegian Veterinary Institute |
| July | 2023 | Nordic | Norway | Nord-Trøndelag | Nærøysund | Black-legged kittiwake | *Rissa tridactyla* | *Laridae* | Charadriiformes | Norwegian Veterinary Institute |
| July | 2023 | Nordic | Norway | Finnmark | Vadsø | Black-legged kittiwake | *Rissa tridactyla* | *Laridae* | Charadriiformes | Norwegian Veterinary Institute |
| July | 2023 | Nordic | Norway | Finnmark | Vadsø | Black-legged kittiwake | *Rissa tridactyla* | *Laridae* | Charadriiformes | Norwegian Veterinary Institute |
| July | 2023 | Nordic | Norway | Finnmark | Vardø | Black-legged kittiwake | *Rissa tridactyla* | *Laridae* | Charadriiformes | Norwegian Veterinary Institute |
| July | 2023 | Nordic | Norway | Rogaland | Karmøy | Black-legged kittiwake | *Rissa tridactyla* | *Laridae* | Charadriiformes | Norwegian Veterinary Institute |
| July | 2023 | Nordic | Norway | Finnmark | Vadsø | Black-legged kittiwake | *Rissa tridactyla* | *Laridae* | Charadriiformes | Norwegian Veterinary Institute |
| July | 2023 | Nordic | Norway | Finnmark | Vadsø | Black-legged kittiwake | *Rissa tridactyla* | *Laridae* | Charadriiformes | Norwegian Veterinary Institute |
| July | 2023 | Nordic | Norway | Finnmark | Vadsø | Black-legged kittiwake | *Rissa tridactyla* | *Laridae* | Charadriiformes | Norwegian Veterinary Institute |
| July | 2023 | Nordic | Norway | Finnmark | Vadsø | Black-legged kittiwake | *Rissa tridactyla* | *Laridae* | Charadriiformes | Norwegian Veterinary Institute |
| July | 2023 | Nordic | Norway | Finnmark | Vadsø | Black-legged kittiwake | *Rissa tridactyla* | *Laridae* | Charadriiformes | Norwegian Veterinary Institute |
| July | 2023 | Nordic | Norway | Finnmark | Vadsø | Black-legged kittiwake | *Rissa tridactyla* | *Laridae* | Charadriiformes | Norwegian Veterinary Institute |
| July | 2023 | Nordic | Norway | Finnmark | Vadsø | Black-legged kittiwake | *Rissa tridactyla* | *Laridae* | Charadriiformes | Norwegian Veterinary Institute |
| July | 2023 | Nordic | Norway | Finnmark | Vadsø | Black-legged kittiwake | *Rissa tridactyla* | *Laridae* | Charadriiformes | Norwegian Veterinary Institute |
| July | 2023 | Nordic | Norway | Finnmark | Vadsø | Black-legged kittiwake | *Rissa tridactyla* | *Laridae* | Charadriiformes | Norwegian Veterinary Institute |
| July | 2023 | Nordic | Norway | Finnmark | Vadsø | Black-legged kittiwake | *Rissa tridactyla* | *Laridae* | Charadriiformes | Norwegian Veterinary Institute |

| Submitting lab | Authors |
|---|---|
| Finnish Food Authority | Kauppinen, Ari; Kareinen, Lauri; Tammiranta, Niina; Kantala, Tuija; Fusaro, Alice; Giussani, Edoardo; Gadd, Tuija |
| Finnish Food Authority | Kauppinen, Ari; Kareinen, Lauri; Tammiranta, Niina; Kantala, Tuija; Fusaro, Alice; Giussani, Edoardo; Gadd, Tuija |
| Finnish Food Authority | Kauppinen, Ari; Kareinen, Lauri; Tammiranta, Niina; Kantala, Tuija; Fusaro, Alice; Galiano, Monica; Ikonen, Niina; Lindh, Erika; Gadd, Tuija |
| Statens Serum Institute | Hjulsager,Charlotte; Liang,Yuan |
| Erasmus Medical Center | Not stated |
| Erasmus Medical Center | Bellido, Beatriz |
| Erasmus Medical Center | Not stated |
| Animal and Plant Health Agency (APHA) | Not stated |
| Animal and Plant Health Agency (APHA) | Not stated |
| Istituto Zooprofilattico Sperimentale Delle Venezie | Ruano,M.J.; Rocha,A.; Sanchez,A.; Aguero,M.; Pastori, A.; Zecchin, B.; Fusaro, A.; Schivo, A.; Salviato, A.; Palumbo, E.; Giussani, E.; Monne, I.; Terregino, C. |
| Austrian Agency for Health and Food Safety (AGES) | Not stated |
| National Veterinary Research Institut Poland, PIWet-PIB | Swieton,Edyta; Tarasiuk,Karolina; Wyrostek,Krzysztof; Kozak,Edyta; Stys-Fijol,Natalia; Smietanka,Krzysztof |
| State Veterinary Institute Prague | Alexander,Nagy; Lenka,Cernikova; Martina,Stara |
| Sciensano, Department of Animal Infectious Diseases | Van Borm, Steven; Vandenbussche, Frank; Roupie, Virginie; Lambrecht, Benedicte; Van Borm, Steven; Vandenbussche, Frank; Roupie, Virginie; Lambrecht, Benedicte; Steensels, Mieke |
| ANSES Agence Nationale De Securite Sanitaire De L'alim | Not stated |
| State Research Center of Virology and Biotechnology (VEC | Natalia,Goncharova; Ivan,Susloparov; Natalia,Kolosova; Alexey,Danilenko; Juliya,Bulanovich; Vasiliy,Marchenko; Alexander,Ryzhikov |
| Norwegian Veterinary Institute | Gjerset, Britt; Bøe, Cathrine A. |
| Norwegian Veterinary Institute | Gjerset, Britt; Bøe, Cathrine A. |
| National Veterinary Research Institut Poland, PIWet-PIB | Swieton, E.; Domanska-Blicharz, K.; Tarasiuk, K.; Wyrostek, K.; Stys-Fijol, N. |
| Finnish Food Authority | Kauppinen, Ari; Kareinen, Lauri; Tammiranta, Niina; Kantala, Tuija; Fusaro, Alice; Gadd, Tuija |
| Istituto Zooprofilattico Sperimentale Delle Venezie | Ruano,M.J.; Rocha,A.; Sanchez,A.; Aguero,M.; Pastori, A.; Zecchin, B.; Fusaro, A.; Schivo, A.; Salviato, A.; Palumbo, E.; Giussani, E.; Monne, I.; Terregino, C. |
| Swedish Veterinary Agency (SVA) | Siamak,Zohari |
| Istituto Zooprofilattico Sperimentale Delle Venezie | Krivko, L.; Pastori, A.; Zecchin, B.; Fusaro, A.; Schivo, A.; Salviato, A.; Palumbo, E.; Giussani, E.; Monne, I.; Terregino, C. |
| Norwegian Veterinary Institute | Gjerset, Britt; Bøe, Cathrine A. |
| Norwegian Veterinary Institute | Gjerset, Britt; Bøe, Cathrine A. |
| Norwegian Veterinary Institute | Gjerset, Britt; Bøe, Cathrine A. |
| Norwegian Veterinary Institute | Gjerset, Britt; Bøe, Cathrine A. |
| Norwegian Veterinary Institute | Gjerset, Britt; Bøe, Cathrine A. |
| Norwegian Veterinary Institute | Gjerset, Britt; Bøe, Cathrine A. |
| Norwegian Veterinary Institute | Gjerset, Britt; Bøe, Cathrine A. |
| Norwegian Veterinary Institute | Gjerset, Britt; Bøe, Cathrine A. |
| Norwegian Veterinary Institute | Gjerset, Britt; Bøe, Cathrine A. |
| Norwegian Veterinary Institute | Gjerset, Britt; Bøe, Cathrine A. |
| Norwegian Veterinary Institute | Gjerset, Britt; Bøe, Cathrine A. |
| Norwegian Veterinary Institute | Gjerset, Britt; Bøe, Cathrine A. |
| Norwegian Veterinary Institute | Gjerset, Britt; Bøe, Cathrine A. |
| Norwegian Veterinary Institute | Gjerset, Britt; Bøe, Cathrine A. |
| Norwegian Veterinary Institute | Gjerset, Britt; Bøe, Cathrine A. |
| Norwegian Veterinary Institute | Gjerset, Britt; Bøe, Cathrine A. |
| Norwegian Veterinary Institute | Gjerset, Britt; Bøe, Cathrine A. |
| Norwegian Veterinary Institute | Gjerset, Britt; Bøe, Cathrine A. |
| Norwegian Veterinary Institute | Gjerset, Britt; Bøe, Cathrine A. |
| Norwegian Veterinary Institute | Gjerset, Britt; Bøe, Cathrine A. |
| Norwegian Veterinary Institute | Gjerset, Britt; Bøe, Cathrine A. |
| Norwegian Veterinary Institute | Rødmo, Grim; Gjerset, Britt; Bøe, Cathrine A. |
| Norwegian Veterinary Institute | Rødmo, Grim; Gjerset, Britt; Bøe, Cathrine A. |
| Norwegian Veterinary Institute | Rødmo, Grim; Gjerset, Britt; Bøe, Cathrine A. |
| Norwegian Veterinary Institute | Rødmo, Grim; Gjerset, Britt; Bøe, Cathrine A. |
| Norwegian Veterinary Institute | Rødmo, Grim; Gjerset, Britt; Bøe, Cathrine A. |
| Norwegian Veterinary Institute | Rødmo, Grim; Gjerset, Britt; Bøe, Cathrine A. |
| Norwegian Veterinary Institute | Rødmo, Grim; Gjerset, Britt; Bøe, Cathrine A. |
| Norwegian Veterinary Institute | Rødmo, Grim; Gjerset, Britt; Bøe, Cathrine A. |
| Norwegian Veterinary Institute | Rødmo, Grim; Gjerset, Britt; Bøe, Cathrine A. |
| Norwegian Veterinary Institute | Rødmo, Grim; Gjerset, Britt; Bøe, Cathrine A. |
