## Appendix 3 for "Highly Pathogenic Avian Influenza A(H5N1) Caused Mass Death among Black-legged Kittiwakes (*Rissa tridactyla*) in Norway, 2023"

**Appendix 3. Table 1. Markers which might indicate mammalian adaptation identified using FluMut* in whole genome sequenced HPAI H5N1 clade 2.3.4.4b genotype BB viruses from 10 black-legged kittiwakes at Ekkerøy in Norway, 2023.**

| Protein | Marker | Effect | Subtype | References |
|---|---|---|---|---|
| PB2 | L89V, G309D | Increased polymerase activity in mammalian cells, Increased virulence in mice | H5N1 | Li J. et al., 2009; Suttie A. et al., 2019 |
| PB2 | K389R | Increased polymerase activity in mammalian cells, Increased replication in mammalian cells | H7N9 | Hu M. et al., 2017b; Suttie A. et al., 2019 |
| PB2 | L89V, G309D, T339K, R477G, I495V, K627E, A676T | Increased polymerase activity in mammalian cells, Increased virulence in mice | H5N1 | Li J. et al., 2009; Suttie A. et al., 2019 |
| PB2 | V598T | Increased replication in mammalian cells, Increased polymerase activity in mammalian cells, Increased virulence in mice | H7N9 | Hu M. et al., 2017b; Suttie A. et al., 2019 |
| PB2 | K699R | Enhanced virulence in mice, Increased viral replication in mammalian cells | H1N1 | Zhang T. et al., 2017 |
| PB1 | D3V | Increased polymerase activity and replication in avian and mammalian cells | H5N1 | Elgendy E. et al., 2017; Suttie A. et al., 2019 |
| PB1 | D622G | Increased polymerase activity in mammalian cells, increased polymerase activity and virulence in mice | H5N1 | Feng X. et al., 2016; Suttie A. et al., 2019 |
| PB1-F2 | N66S | Enhanced antiviral response, replication and virulence in mice | H5N1 | Conenello G. et al., 2007; Schmolke M. et al., 2011; Suttie A. et al., 2019 |
| PA | S37A | Increased polymerase activity in mammalian cells | H7N9 | Suttie A. et al., 2019; Yamayoshi S. et al., 2014 |
| PA | N383D | Increased polymerase activity in avian and mammalian cells | H5N1 | Song J. et al., 2011; Song J. et al., 2015; Suttie A. et al., 2019 |
| PA | N409S | Increased polymerase activity and replication in mammalian cells | H7N9 | Suttie A. et al., 2019; Yamayoshi S. et al., 2014 |
| HA1-5 | S107R, T108I | Increased pH of fusion, increased virulence in chickens and mice | H5N1 | Suttie A. et al., 2019; Wessels U. et al., 2018 |
| HA1-5 | S133A | Increased pseudovirus binding to α2-6 | H5N1 | Suttie A. et al., 2019; Yang Z. et al., 2007 |
| HA1-5 | T134A | Increased viral replication in mice lungs, Increased virus thermostability | H9N2 | Zhang J. et al., 2023 |
| HA1-5 | S154N | Increased virus binding to α2-6 | H5N1 | Suttie A. et al., 2019; Wang W. et al., 2010 |
| HA1-5 | T156A | Increased transmission in guinea pigs, Increased virus binding to α2-6 | H5N1 | Gao Y. et al., 2009; Suttie A. et al., 2019; Wang W. et al., 2010 |
| HA1-5 | V182N | Decreased virus binding to α2-3, Increased virus binding to α2-6 | H13N6 | Lu X. et al., 2013; Suttie A. et al., 2019 |
| HA1-5 | K218Q, S223R | Increased virus binding to α2-3 and α2-6 | H5N1 | Guo H. et al., 2017; Suttie A. et al., 2019 |
| HA2-5 | K64E | Decreased HA stability, Decreased virulence in mice, Increased pH of fusion | H7N9 | Sun X. et al., 2019; Suttie A. et al., 2019 |
| NA | A369I | Distruption of the second sialic acid binding site (2SBS) | H5N1, Unknown | de Vries E. et al., 2023, Du W. et al., 2018 |
| NP | Y52N | Evade human BTN3A3 (inhibitor of avian influenza A viruses replication) | Unknown | Pinto R. et al., 2023 |
| M2 | A30S | Increased resistance to amantadine and rimantadine | H5N1, H5N2, H7N2 | Bean W. et al., 1989; Cheung C. et al., 2006; Ilyushina N. et al., 2005; Suttie A. et al., 2019 |
| NS1 | P42S | Decreased antiviral response, and increased virulence in mice | H5N1 | Jiao P. et al., 2008; Suttie A. et al., 2019 |
| NS1 | L103F, I106M | Increased virulence in mice | H5N1 | Kuo R. et al., 2009; Spesock A. et al., 2011; Suttie A. et al., 2019 |
| NS1 | I106M | Increased viral replication in mammalian cells, Increased virulence in mice | H1N1 with all internal genes from H7N9 | Ayllon J. et al., 2014; Suttie A. et al., 2019 |
| NS1 | C138F | Decreased interferon response, Increased viral replication in mammalian cells | H5N1 | Li J. et al., 2018; Suttie A. et al., 2019 |
| NS1** | K55E, K66E, C138F | Decreased interferon response, Enhanced replication in mammalian cells | H5N1 | Li J. et al., 2018; Suttie A. et al., 2019 |
| NS1, NS2 | NS1-205S, NS-2:T48A | Decreased antiviral response in ferrets | H5N1 | Imai H. et al., 2010; Suttie A. et al., 2019 |

*https://github.com/izsvenezie-virology/FluMut, FluMutGUI 3.1.1; FluMut 0.6.3; FluMutDB 6.3, released on 2024-09-12
**present in all viruses, except one (bird no. 9 K55G, K66E, C138F)
